# Researchers‘ perspectives on preregistration in animal research

**DOI:** 10.1101/2025.11.07.687141

**Authors:** Cristina Priboi, Boris Mayer, Evie Vergauwe, Bernice Elger, Hanno Würbel

**Affiliations:** Veterinary Public Health Institute, University of Bern, Bern, Switzerland; Institute of Psychology, University of Bern, Switzerland; Faculty of Psychology and Educational Sciences, University of Geneva, Switzerland; Institute for Biomedical Ethics, University of Basel, Switzerland

## Abstract

Preregistration is arguably one of the most promising and impactful Open Science practices. Defined as the a priori registration of study designs and analysis plans, preregistration has long been established as standard practice in clinical human research and is increasingly taken up in other fields of science. Despite growing evidence suggesting that preregistration can mitigate questionable research practices, and the existence of two platforms targeting animal studies, preregistration remains uncommon in animal research. In light of the reproducibility crisis and calls for more transparency and rigor also in animal research, preregistration represents a potentially promising step forward. However, implementing such policies without understanding their impact carries potential risks. It is, therefore, essential to uncover the strengths, weaknesses, opportunities, and threats of preregistration before advancing its implementation in animal research. This current study addressed this need as part of a larger feasibility project on preregistration of animal experiments in Switzerland, and aimed to: 1) assess the researchers experiences with preregistration; 2) examine their attitudes, subjective norms, perceived behavioral control, intentions, motivations, and perceived obstacles regarding preregistration; 3) explore associations between these psychosocial constructs and relevant background characteristics; 4) identify perceived facilitators and barriers to preregistration; and 5) summarize researchers’ suggestions for improving preregistration. A preregistered cross-sectional online survey was conducted among all registered study directors of ongoing animal experiments in Switzerland. Of the 1,385 invited study directors, 418 completed the survey (30.2% return rate; 41% female; age *M* = 47.1, *SD* = 9.52). Among them, 39.2% had never heard of preregistration, and only 10% had preregistered studies before participating in the survey. Bureaucratic burden (77.6%), time costs (71.4%), and low flexibility (65.7%) were the most common reported barriers to preregistration. On average, participants described rather unfavorable attitudes towards preregistration, negative subjective norms, relatively low perceived behavioral control, weak intention, and limited motivation to preregister, along with high perceived obstacles. Participants who had never preregistered a study, as well as those with more research experience, showed more negative scores on all assessed psychosocial constructs related to preregistration. Our findings offer guidance on promising measures to enhance acceptance of preregistration among animal researchers, including raising awareness, offering education and training, and facilitating procedures, to enhance the acceptance of preregistration among animal researchers.

## Introduction

The scientific community is facing a profound methodological reformation towards Open Science in response to the so-called reproducibility crisis. Institutions, funders and journals are implementing policies to enhance rigor, transparency and reproducibility, with the goal of increasing credibility and trust in science. Preregistration is a promising tool for improving transparency and quality of research, and is arguably among the most impactful Open Science practices. Defined as the registration of hypotheses, study designs and analysis plans in independent open repositories before data collection or analysis (1–4), preregistration may reduce the use of questionable research practices such as publication bias, selective reporting, *p*-hacking, and HARKing, leading to more reproducible research findings (1–3, 5–9). Although the effects of preregistration have only recently begun to be studied systematically, and robust evidence is still scarce, available research from different disciplines provides direct or indirect support for a positive impact of preregistration on the reproducibility of research findings (10, 11). Preregistered studies report lower proportions of statistically significant results, find smaller effect sizes, and provide more thorough methodological descriptions compared to studies that have not been preregistered (5, 8, 12–17).

The inability to replicate published results, e.g. in cancer research (18), and the low rate of translating preclinical findings to clinical studies (19, 20) have questioned the rigor and reproducibility of animal research (18, 21–23). This could reduce public support for the responsible use of animals in research, which depends not only on the implementation of the 3Rs but also on the assumption that such practices will lead to important contributions to science and medical progress, or nature conservation. It has been shown that study protocols and results from many research projects using animals are never published or not published in full (24–27). Furthermore, substantial deficiencies in the use and reporting of measures against risks of bias have been documented (28, 29), including in an exemplary study conducted in Switzerland (30, 31). Apart from increasing public doubts about animal research, these questionable research practices can slow down scientific progress, waist animal lives for inconclusive results, and expose participants in clinical trials to unnecessary risks (32–34).

Although study registries tailored to animal research have been established (1, 4), the animal research community appears to meet preregistration with reservations (34). This may be due to a number of reasons, including researchers’ concerns that preregistration would increase administrative burden, that it would stifle creativity and discovery, that others might steal their ideas, or that they would become targets of animal rights activists (17, 35–37). In addition, researchers may not be aware of the possibilities, functioning, and benefits of preregistration (34). Nevertheless, policies regarding preregistration are already shifting with key stakeholders adopting preregistration standards for animal studies. A report by the US National Institute of Health working group has recommended preregistration for animal studies (38), an increasing number of journals are adopting preregistration badges as part of their publication policies (3, 8), and a recent call has been made to implement preclinical study preregistration in the animal research community (37). Despite the growing popularity of preregistration, the relevance, applicability, and implementation of preregistration have not yet been properly analyzed for animal research. There is a risk of changing policies too far or prematurely before the potential costs and benefits for innovation and research quality, and the practical implications of preregistration have been understood by all parties involved. There is, therefore, an urgent need to evaluate the perceived strengths, weaknesses, opportunities, and threats of preregistration in animal research. As part of a project to address these different aspects, we conducted the present survey among all animal study directors registered in Switzerland. The study was informed by Spitzer and Mueller (39), who investigated attitudes and experiences with preregistration in psychology researchers. Building on their framework, the specific aims of our study were to:

1. Assess the experience of animal researchers in Switzerland with study preregistration.
2. Examine several psychosocial constructs related to preregistration, including attitudes, subjective norms, perceived behavioral control, intentions, motivations, and perceived obstacles.
3. Explore associations between these six psychosocial constructs and background characteristics, such as animal researchers’ preregistration experience, animal research experience, gender, field of animal research, and organization of employment.
4. Identify perceived barriers and facilitators to study preregistration from the perspective of Swiss animal researchers.
5. Summarize participants’ suggestions for improving the practice of preregistration in Switzerland.

We reasoned that achieving these aims is important to provide guidance on promising measures to enhance acceptance of preregistration among animal researchers and facilitate its implementation.

## Materials and methods

The aims, design and analysis plan of this study were preregistered during data collection on the Open Science Framework on June 25, 2024 (https://doi.org/10.17605/OSF.IO/CAUFW). All deviations from the preregistered plan are described following Willroth and Atherton (40) and can be found in the supplementary material (S1 Table).

### Study design

This cross-sectional study received ethical approval from the Ethical Commission of the Faculty of Human Sciences at the University of Bern, Switzerland (Nr. 2024-03-06).

### Sample

The study sample included researchers accredited as “study directors” of animal experiments under Swiss law. In Switzerland, all applications for animal experiments must be submitted through the federal platform Animex-ch. According to Swiss animal welfare legislation (TSchV Art. 131), study directors are responsible for designing and conducting animal experiments in compliance with the national and cantonal legal requirements. To qualify as a study director, researchers need to hold a biomedical university degree, have at least three years of experience with animal experimentation, and complete specific additional education and training. Importantly, the role of “study director” is a legal designation and does not necessary overlap with the role of “principal investigator”. Participants of this study were sampled from this population of study directors. Only researchers who held an active animal research license and were registered as study directors or deputy study directors at the time of recruitment were eligible to participate.

### Procedure & data collection

Data collection was conducted with the support of the Federal Food Safety and Veterinary Office (FSVO) in Switzerland and took place between 29.05.2024 and 26.06.2024. The FSVO, which holds the contact information for all registered study directors and deputy study directors in the country, sent an official invitation email with the survey link to all eligible study directors. Two further reminders were sent by the FSVO, seven and 14 days after the initial invitation email. Additionally, one more reminder was issued at the institutional level by the Animal Welfare Officers (AWO). At the time of the study, 49 institutions in Switzerland had designated AWOs, whose role, as defined by the Animal Welfare Ordinance, is to ensure that applications for animal experiments are complete and coherent, particularly regarding the justification of the experiment’s necessity, defined endpoints and harm-reduction measures, and the harm–benefit analysis.

### Questionnaires

Participants completed an online survey in English that included both closed-ended and open-ended items. The items were adapted from the questionnaire developed by Spitzer and Mueller (39) and modified to align with the field of animal research and the study’s research questions. An overview of the specific changes can be consulted in the supplementary material (S2 Codebook). The online survey collected information related to study preregistration, such as participants’ experience with preregistration, their attitudes, subjective norms, perceived behavioral control, intentions, motivations, obstacles, facilitators, barriers, and suggestions for improving preregistration, as well as socio-demographic characteristics.

#### Socio-demographic characteristics

The following demographic variables were collected from the sample: age, gender, years registered as study directors, years of animal research experience, academic age, educational attainment, seniority level, organization of employment, and field of animal research.

#### General experiences

Between three and seven closed-ended items, depending on whether participants had ever preregistered a study or not, were used to assess participants’ general practices and their experiences with study preregistration (S2 Codebook).

#### Psychosocial constructs

The survey included six Preregistration Scales, each evaluating one of the following six psychosocial constructs related to preregistration: attitudes (23 items); subjective norms (7 items); perceived behavioral control (7 items); intentions (3 items); motivations (10 items); obstacles (10 items). The attitudes scale measured participants’ evaluations of and opinions toward the concept of preregistration. The subjective norms scale explored the social pressure and expectations perceived by participants regarding preregistering their studies. The perceived behavioral control scale assessed participants’ judgment of their own ability and confidence to preregister studies. The intentions scale evaluated the determination to preregister future studies, while the motivations scale focused on understanding the rationales behind researchers’ decisions to preregister or not preregister their studies. The obstacles scale measured challenges perceived by researchers in the preregistration process. A detailed description of the items for each scale is provided in the supplementary material (S2 Codebook). All items were closed-ended and to be responded to on a 7-point Likert scale ranging from 1 to 7. The values were recoded from the original scale to a scale centered around 0 ranging from −3 to +3.

#### Barriers and facilitators

The number and wording of items addressing barriers and facilitators of preregistration varied depending on whether participants had previously preregistered a study or not (S2 Codebook). Barriers were assessed with one closed-ended item and up to four open-ended items *(“What do you perceive as drawbacks of preregistration?”; “What do you think would be the long-term negative consequences of mandatory preregistration?”; “What are the reasons for not preregistering your studies?”; “I am now less motivated to preregister than I was before because_______”)*. Facilitators were examined exclusively with open-ended items (*“What do you perceive as benefits of preregistration?”; “What do you think would be the long-term benefits of mandatory preregistration?”; “I am now more motivated to preregister than I was before because_______”*).

#### Suggestions

The following open-ended items were used to collect suggestions for improving preregistration of animal experiments in Switzerland: *“We are interested in how we can improve various aspects of preregistration (e.g. templates, repositories, reviewing process, integrations in published articles, education etc.). Do you have any suggestions? What do you think should be improved about preregistration?”; “What would make you (and perhaps other researchers) preregister more often?”; “Do you have any suggestions as to how to lower your (or other researchers’) negative perceptions of preregistration?”*.

### Data analysis

#### Sample description

Descriptive statistics are reported to summarize the socio-demographic characteristics of the study participants. To examine differences between participants and non-participants, independent samples *t*-tests were conducted for continuous variables and chi-square tests for categorical variables. The following information was provided by the FSVO: the total number of eligible study directors invited to participate, and, separately for participants and non-participants, their age (range, median, mean, and standard deviation), gender (number and percentage of female and male participants), and years since their registration as study directors (range, median, mean, and standard deviation). Due to differences in the operationalization of the gender variable and to ensure comparability with FSVO data, the survey responses were recoded prior to analysis. The FSVO used two categories (“female” and “male”), while our survey included four options (“female”, “male”, “other”, and “prefer not to say”). For consistency, responses coded as “other” and “prefer not to say” were treated as missing in the comparative analysis. The values made available by the FSVO enabled comparisons between participants and non-participants with respect to age, gender, and years since registration as study directors.

#### General experiences (Aim 1)

Descriptive statistics were calculated and reported for each item assessing participants’ general experiences with preregistration. In addition, participants with and without preregistration experience were compared across all socio-demographic characteristics. Independent samples *t*-tests were used to analyze mean differences in continuous variables, while percentage differences in categorical variables were tested using Fisher’s Exact Test.

#### Psychosocial constructs (Aim 2)

##### Psychometric quality

The items of the scales used in the survey were adapted from a questionnaire developed by Spitzer and Mueller (39), who reported high reliabilities for all six Preregistration Scales. However, it remained unclear whether these scales were truly unidimensional. Therefore, we performed principal component analyses (PCA) to explore the dimensional structure of the items within each of the six scales. Subsequently, we calculated McDonald’s omega to assess the internal consistency of each scale (or subscale–in case of more than one dimension obtained in the PCA for a specific scale). To determine the appropriate number of components, we applied both the scree-plot criterion and parallel analysis, using the mean eigenvalues of 1,000 randomly generated samples for comparison. If more than one component was identified for a specific scale, the extraction was followed by an oblique Oblimin-rotation to arrive at a simple structure even when components were correlated.

##### Overview scales

Descriptive results are reported for each of the six Preregistration Scales (attitudes, subjective norms, perceived behavioral control, intentions, motivations, obstacles).

#### Association analysis (Aim 3)

A multivariate analysis of covariance (MANCOVA) was conducted to examine the overall effect of several predictors on all Preregistration Scales, which served as the outcome variables. The predictors included in the multivariate model were: preregistration experience (“preregistered at least one study”, “never preregister a study”), animal research experience (in years), gender (“female”, “male”, “prefer not to say”), field of animal research (“basic biological research”, “general biology”, “basic and experimental medical research”), and organization of employment (“academia”, “private sector”, “other”). Due to the very low frequency of responses in the “other” gender category, this category was recoded as missing prior to analysis. Wilks’ Lambda and Pillai’s Trace were used to test the overall effects of the MANCOVA model for each predictor variable. If a significant multivariate effect was detected, univariate analyses of covariance (ANCOVAs) were performed for each outcome separately to identify which predictors were significantly associated with which outcome. Partial eta-squared (*η*_*p*_^2^) was computed for each predictor within each outcome to estimate the effect sizes in the univariate models. Values of 0.01, 0.06, and 0.14 for *η*_*p*_^2^ were considered small, medium and large (41). For categorical predictors with more than two levels which showed significant effects in the ANCOVAs, post hoc pairwise comparisons were conducted using Tukey’s correction for multiple comparisons.

#### Thematic analysis (Aims 4 and 5)

The answers provided to the qualitative open-ended items were analyzed using thematic analysis (42, 43). The data were inductively coded and categorized into facilitators, barriers, and suggestions for improving preregistration. In a subsequent step, recurring themes were identified and described within each category.

## Results

### Sample description

The FSVO identified 1,428 eligible study directors and invited them via email to take part in the online survey. Each study director was provided with a unique access code required to enter the survey. Forty-three emails bounced back, resulting in 1,385 successfully reached study directors. Of these, 744 individuals clicked on the survey link. However, 58 did not enter an access code, 59 entered an incorrect code, and 106 either failed to check the consent box or explicitly declined consent, preventing them from proceeding. Additionally, 17 access codes were used to complete the questionnaire twice, and 2 codes were used three times. In such cases, only the first completed attempt was kept, and subsequent entries were deleted. Three additional cases were removed at the request of the participants. After these exclusions, 497 valid entries remained. A further 17 cases were excluded due to over 90% missing data, and 62 for failing the attention check item (S3 Image). A final sample of *N* = 418 participants was included in the analysis, resulting in a response rate of 30.2% among the reached population.

#### Socio-demographic characteristics

Participants had a mean age of 47.1 (*SD* = 9.52) years, and slightly less than half identified as female (Table 1). On average, participants were registered as study directors for 12.6 (*SD* = 5.86) years and worked in animal research for 20.1 (*SD* = 8.83) years. The majority were highly educated, with over 90% reporting a PhD, Dr. med. or higher, and 60% occupying senior positions such as group leader or professor. The predominant research field was basic and experimental medical research (72.6%), and most participants were employed in academic institutions (77.6%).

**Table 1:**
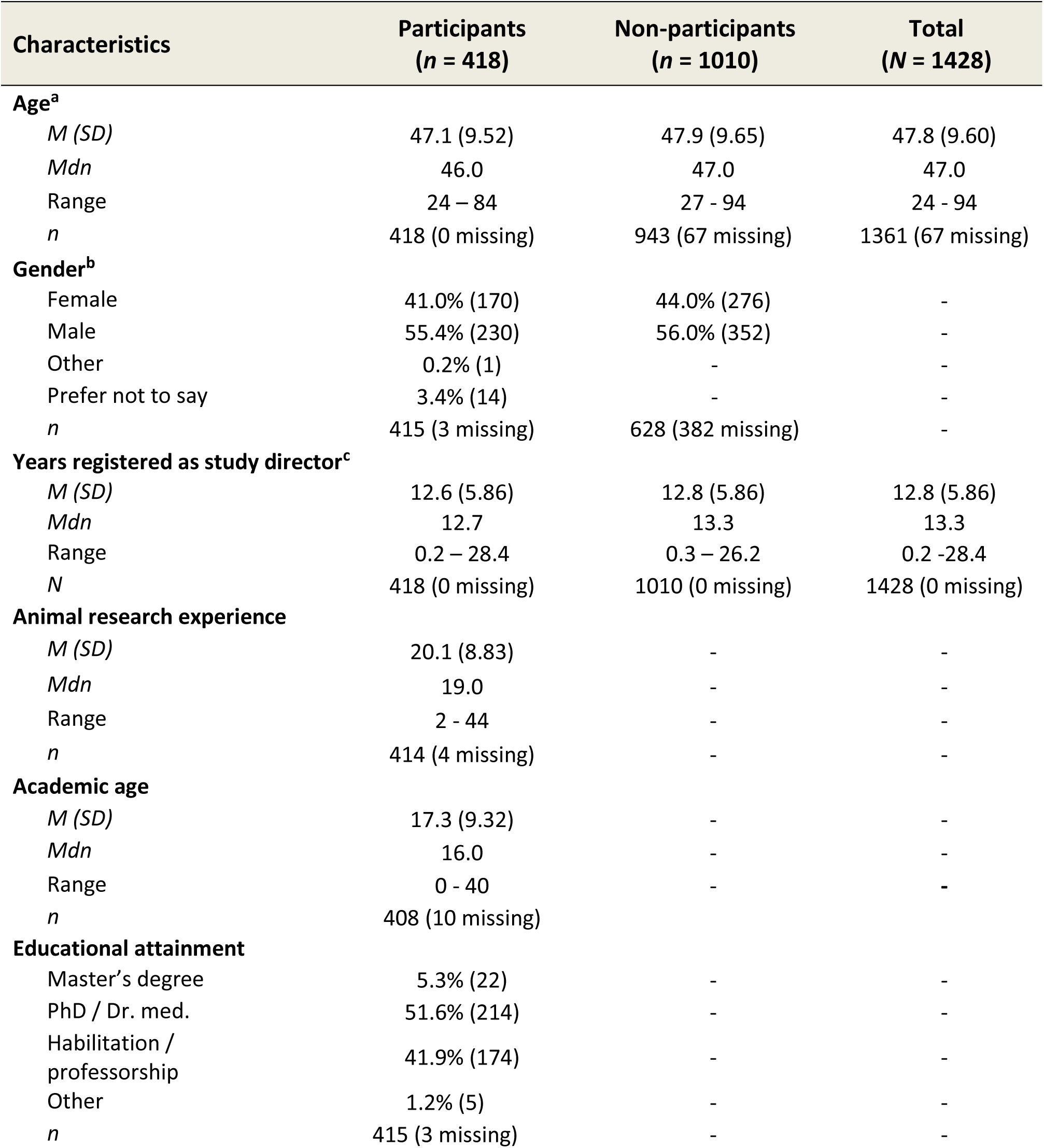
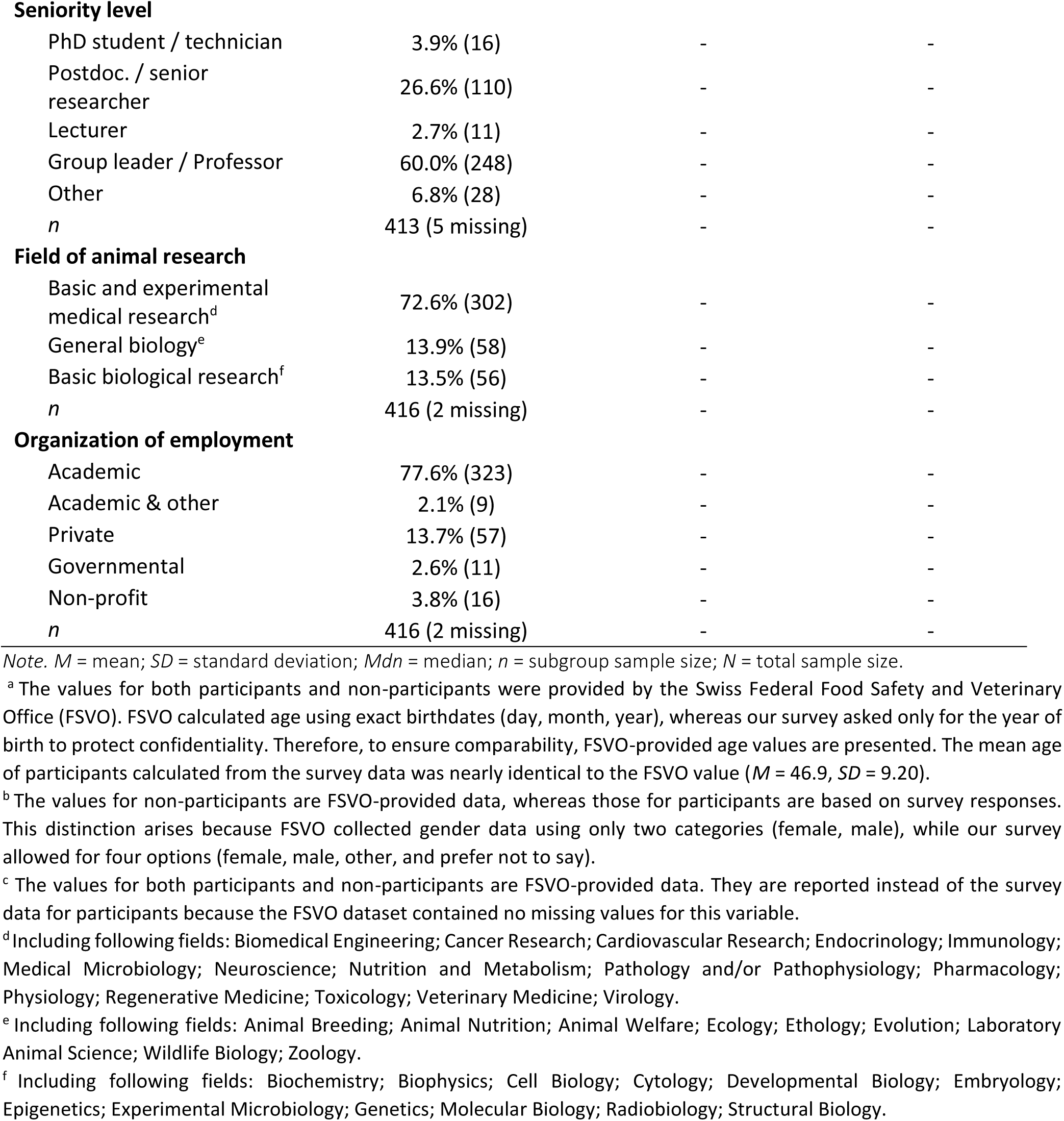
Sample description.

#### Comparison participants vs. non-participants

An independent samples *t*-test was conducted to compare the mean age of participating study directors (*M* = 47.1, *SD* = 9.52, *N* = 418) with that of non-participating study directors (*M* = 47.9, *SD* = 9.65, *N* = 1010). The analysis revealed no statistically significant difference between the two groups, *t*(809.2) = −1.59, *p* = .112. A chi-square test of independence was also conducted to examine whether gender distribution differed between participants and non-participants. The association was not statistically significant, *χ*²(1, *N* = 1028) = 0.22, *p* = .642, suggesting that the proportion of male and female study directors was similar across the two group (participants: 42.5% female and 57.5% male; non-participants: 44% female and 56% male). Similarly, the average number of years registered as study director did not significantly differ between participants (*M* = 12.6, *SD* = 5.85, *N* = 418) and non-participants (*M* = 12.8, *SD* = 5.85, *N* = 943), *t*(778.8) = −0.50, *p* = .619. Taken together, these results suggest that participants and non-participants were comparable in terms of age, gender distribution, and animal research experience.

### General experiences (Aim 1)

Out of the total sample, 10.0% (*n* = 42) reported having previously preregistered at least one study, while 90.0% (*n* = 376) have never done so (S4 Table). Overall, 39.2% (*n* = 147) of participants had never heard of preregistration prior to completing the online survey, while 31.5% (*n* = 118) learned about it through independent readings, and 27.2% (*n* = 102) were introduced to it during further education (S4 Table). Institutional policies were most frequently chosen as influencing the decision to preregister or not (47.6%, *n* = 175), followed by funding guidelines (35.6%, *n* = 131), superior(s) (32.6%, *n* = 120), and journal guidelines (26.4%, *n* = 97). Notably, 28.0% of participants (*n* = 103) stated that nobody can influence their decision (S4 Table). Among participants with preregistration experience, the median number of preregistered studies was 3.0 (range 1 - 80) and the estimated median time required to preregister a study was 11.0 hours (range 1 - 150) (S4 Table). Furthermore, the comparison of participants with and without preregistration experience revealed no statistically significant differences across all tested socio-demographic characteristics, indicating that the two groups were largely comparable (S5 Table).

### Psychosocial constructs (Aim 2)

#### Psychometric quality

The dimensionality and internal consistency of all preregistration scales was assessed using PCA and McDonald’s omega. When necessary, items with poor psychometric properties and weak conceptual alignment were removed, resulting in improved internal consistency and construct validity of each scale. The Preregistration Scales for Attitudes, Subjective Norms, Intentions, and Motivations showed clear unidimensional structures and good to excellent reliability (ω = .84 to ω = .96). In contrast, the Perceived Behavioral Control Scale and the Obstacles Scale revealed a two-component structure and were therefore each divided into two subscales. The Perceived Behavioral Control Scale was separated into: the Perceived Behavioral Control – Resources Subscale (ω = .72), reflecting resources and support for preregistration from colleagues and supervisors; and the Perceived Behavioral Control – Knowledge Subscale (ω = .65), reflecting the knowledge required and authority to decide about preregistration. Similarly, the Obstacles Scale was structured into: the Practical Obstacles Subscale (ω = .84), reflecting perceived disadvantages in terms of time and effort; and the Competitive Obstacles Subscale (ω = .66), describing concerns about scooping and competitive challenges. For detailed results of these analyses, see S6 Psychometric quality.

#### Overview of scales

Based on the psychometric analysis, the following Preregistration Scales and Subscales were used as outcomes in all subsequent analyses: attitudes scale (21 items), subjective norms scale (6 items), perceived behavioral control – resources subscale (4 items), perceived behavioral control – knowledge subscale (3 items) intentions scale (3 items), motivations scale (10 items), practical obstacles subscale (6 items), and competitive obstacles subscale (4 items).

Descriptive statistics of all Preregistration Scales and Subscales are presented in Table 2. In the total sample, participants reported slightly negative attitudes toward preregistration (*M* = −0.9, *SD* = 1.22), low subjective norms (*M* = −1.6, *SD* = 1.00), and moderate perceived behavioral control (resources subscale: *M* = −0.7, *SD* = 1.06; knowledge subscale: *M* = −0.4, *SD* = 1.30). Similarly, both intentions (*M* = −0.8, *SD* = 1.65) and motivations to preregister (*M* = −0.8, *SD* = 1.36) were rather low. In contrast, perceived obstacles were rated to be rather high (practical obstacles: *M* = 1.5, *SD* = 1.04; competitive obstacles: *M* = 0.7, *SD* = 1.05), indicating that many participants perceived substantial barriers to preregistration.

**Table 2:** Overview Preregistration Scales.

| Scales | Total participants<br>($N = 418$) |
| --- | --- |
| **Attitudes Scale** |  |
| $M (SD)$ | -0.9 (1.22) |
| $Mdn$ | -0.8 |
| Range | [-3.0; +2.2] |
| $N$ | 371 (47 missing) |
| **Subjective Norms Scale** |  |
| $M (SD)$ | -1.6 (1.00) |
| $Mdn$ | -1.7 |
| Range | [-3.0; +1.8] |
| $N$ | 353 (65 missing) |
| **Perceived Behavioral Control** |  |
| *Resources Subscale* |  |
| $M (SD)$ | -0.7 (1.06) |
| *Mdn* | -0.8 |
| Range | [-3.0; +2.2] |
| *N* | 351 (67 missing) |
| *Knowledge Subscale* |  |
| *M (SD)* | -0.4 (1.30) |
| *Mdn* | -0.8 |
| Range | [-3.0; +3.0] |
| *N* | 351 (67 missing) |
| **Intentions Scale** |  |
| *M (SD)* | -0.8 (1.65) |
| *Mdn* | -0.7 |
| Range | [-3.0; +3.0] |
| *N* | 349 (69 missing) |
| **Motivations Scale** |  |
| *M (SD)* | -0.8 (1.36) |
| *Mdn* | -0.8 |
| Range | [-3.0; +2.2] |
| *N* | 348 (70 missing) |
| **Obstacles** |  |
| *Practical Obstacles Subscales* |  |
| *M (SD)* | 1.5 (1.04) |
| *Mdn* | 1.7 |
| Range | [-2.2; +3.0] |
| *N* | 348 (70 missing) |
| *Competitive Obstacles Subscales* |  |
| *M (SD)* | 0.7 (1.05) |
| *Mdn* | 0.8 |
| Range | [-2.5; +3.0] |
| *N* | 348 (70 missing) |

Note. *M* = mean; *SD* = standard deviation; *Mdn* = median; *N* = total sample size.

### Association analysis (Aim 3)

A MANCOVA was conducted to examine the effects of preregistration experience, animal research experience, gender, field of research, and organization of employment on the eight Preregistration Scales and Subscales. The multivariate model revealed significant effects of all predictors: preregistration experience [Wilks’ *Λ* = 0.815, *F*(8, 326) = 9.25, *p* < .001]; animal research experience [Wilks’ *Λ* = 0.935, *F*(8, 326) = 2.84, *p* = .005]; gender [Wilks’ *Λ* = 0.862, *F*(16, 652) = 3.13, *p* = .001]; field of research [Wilks’ *Λ* = 0.891, *F*(16, 652) = 2.42, *p* = .002]; and organization of employment (Wilks’ *Λ* = 0.904, *F*(16, 652) = 2.12, *p* = .007). The Pillai’s Trace test confirmed these results.

#### Preregistration experience

The separate univariate ANCOVAs showed that having preregistration experience was significantly associated with more positive attitudes [*b* = 1.16, *F*(1, 333) = 21.10, *p* < .001, *η*_*p*_^2^ = .06]; higher subjective norms [*b* = 1.10, *F*(1, 333) = 22.56, *p* < .001, *η*_*p*_^2^ = .06]; greater perceived behavioral control regarding both resources [*b* = 0.88, *F*(1, 333) = 13.42, *p* < .001, *η*_*p*_^2^ = .04] and knowledge [*b* = 1.36, *F*(1, 333) = 25.38, *p* < .001, *η*_*p*_^2^ = .07]; higher intentions [*b* = 1.31, *F*(1, 333) = 12.62, *p* < .001, *η*_*p*_^2^ = .04]; higher motivations [*b* = 1.11, *F*(1, 333) = 14.22, *p* < .001, *η*_*p*_^2^ = .04]; and fewer perceived practical obstacles [*b* = −0.79, *F*(1, 333) = 12.44, *p* < .001, *η*_*p*_^2^ = .03]. No significant association was found for perceived competitive obstacles [*b* = −0.37, *F*(1, 333) = 2.26, *p* = .134, *η*_*p*_^2^ = .01].

#### Animal research experience

By contrast, more experience in animal research was negatively associated with the outcomes in the univariate ANCOVAs. Longer research experience was linked to less positive attitudes [*b* = −0.02, *F*(1, 333) = 12.72, *p* < .001, *η*_*p*_^2^ = .04]; lower perceived behavioral control – resources [*b* = −0.01, *F*(1, 333) = 4.42, *p* = .036, *η*_*p*_^2^ = .01]; lower intentions [*b* = −0.03, *F*(1, 333) = 10.43, *p* = .001, *η*_*p*_^2^ = .03]; and lower motivations to preregister [*b* = −0.03, *F*(1, 333) = 15.17, *p* < .001, *η*_*p*_^2^ = .04]. In addition, animal research experience showed no significant effects for subjective norms [*b* = −0.001, *F*(1, 333) = 0.14, *p* = .708, *η*_*p*_^2^ = .00]; perceived behavioral control – knowledge [*b* = 0.01, *F*(1, 333) = 0.42, *p* = .520, *η*_*p*_^2^ = .00]; and both practical [*b* = 0.01, *F*(1, 333) = 3.86, *p* = .050, *η*_*p*_^2^ = .01], and competitive obstacles [*b* = −0.005, *F*(1, 333) = 0.19, *p* = .660, *η*_*p*_^2^ = .00].

#### Gender

The univariate analyses also showed differences across genders for attitudes [*F*(2, 333) = 4.99, *p* = .007, *η*_*p*_^2^ = .03]; perceived behavioral control – knowledge [*F*(2, 333) = 16.35, *p* < .001, *η*_*p*_^2^ = .08]; intentions [*F*(2, 333) = 5.79, *p* = .003, *η*_*p*_^2^ = .03]; motivations [*F*(2, 333) = 7.80, *p* < .001, *η*_*p*_^2^ = .04]; and practical obstacles [*F*(2, 333) = 5.20, *p* = .006, *η*_*p*_^2^ = .02]. No significant gender effects were found for subjective norms [*F*(2, 333) = 0.39, *p* = .678, *η*_*p*_^2^ = .00]; perceived behavioral control – resources [*F*(2, 333) = 0.05, *p* = .949, *η*_*p*_^2^ = .00]; and competitive obstacles [*F*(2, 333) = 2.52, *p* = .082, *η*_*p*_^2^ = .01].

Follow-up pairwise comparisons (S7 Table) revealed that participants who preferred not to state their gender reported significantly less positive attitudes and lower motivations to preregister than participants identifying as female or male; also, they reported significantly lower intentions to preregister than those identifying as female. In contrast, females rated their perceived behavioral control – knowledge significantly lower than males as well as those who preferred not to disclose their gender. Although the overall effect of gender on the practical obstacles subscale was significant, no significant differences resulted in the pairwise comparisons of the three gender categories after Tukey correction.

#### Field of research

Differences across animal research fields were found in the univariate ANCOVA models for attitudes [*F*(2, 333) = 5.43, *p* = .005, *η*_*p*_^2^ = .02]; perceived behavioral control – resources [*F*(2, 333) = 3.96, *p* = .020, *η*_*p*_^2^ = .03]; motivations [*F*(2, 333) = 7.02, *p* = .001, *η*_*p*_^2^ = .04]; and competitive obstacles [*F*(2, 333) = 6.06, *p* = .003, *η*_*p*_^2^ = .03]. No significant effects were observed for subjective norms [*F*(2, 333) = 0.13, *p* = .880, *η*_*p*_^2^ = .00]; perceived behavioral control – knowledge [*F*(2, 333) = 0.43, *p* = .653, *η*_*p*_^2^ = .005]; intentions [*F*(2, 333) = 2.60, *p* = .075, *η*_*p*_^2^ = .02]; and for practical obstacles [*F*(2, 333) = 1.86, *p* = .157, *η*_*p*_^2^ = .01].

The subsequent pairwise comparisons (S8 Table) showed that participants working in General Biology reported significantly more positive attitudes towards preregistration, higher motivation, and fewer competitive obstacles compared to those working in Basic Biological Research. They also scored significantly higher on perceived behavioral control – resources and reported fewer competitive obstacles than participants working in Basic and Experimental Medical Research. No further pairwise comparisons between research fields were statistically significant.

#### Organization of employment

The only significant difference across organizations of employment was found for perceived behavioral control – knowledge [*F*(2, 333) = 9.15, *p* < .001, *η*_*p*_^2^ = .05]. No significant effects were observed for attitudes [*F*(2, 333) = 2.40, *p* = .092, *η*_*p*_^2^ = .01]; subjective norms [*F*(2, 333) = 0.19, *p* = .826, *η*_*p*_^2^ = .00]; perceived behavioral control – resources [*F*(2, 333) = 0.04, *p* = .962, *η*_*p*_^2^ = .00]; intentions [*F*(2, 333) = 0.45, *p* = .637, *η*_*p*_^2^ = .00]; motivations [*F*(2, 333) = 0.48, *p* = .619, *η*_*p*_^2^ = .00]; practical obstacles [*F*(2, 333) = 0.24, *p* = .784, *η*_*p*_^2^ = .00]; or competitive obstacles [*F*(2, 333) = 0.51, *p* = .602, *η*_*p*_^2^ = .00].

The pairwise comparisons (S9 Table) revealed that participants employed in academic institutions reported significantly higher perceived behavioral control – knowledge compared to those working in the private industry. No further significant differences between types of organizations of employment were detected.

### Barriers and facilitators (Aim 4)

#### Barriers

Based on the closed-ended items, participants who had never preregistered a study chose bureaucratic burden (77.6%), time costs (71.4%), and low flexibility (65.7%) as the most common barriers to preregistration (Fig 1). Additionally, more than half also indicated the fear of being scooped (53.8%) and the inability to make necessary future changes to their study protocol (52.4%) as important concerns (Fig 1). Similarly, bureaucratic burden (60%), the time-consuming nature of the process (40%), and the insecurity about what to include in the preregistration (33.3%) were the problems most frequent encountered by those who had previously preregistered studies (Fig 1). Further barriers chosen by those with preregistration experience were limited flexibility during analysis (26.7%) and the prevention of the necessary changes of the protocol (20%) (Fig 1).

**Figure 1:**
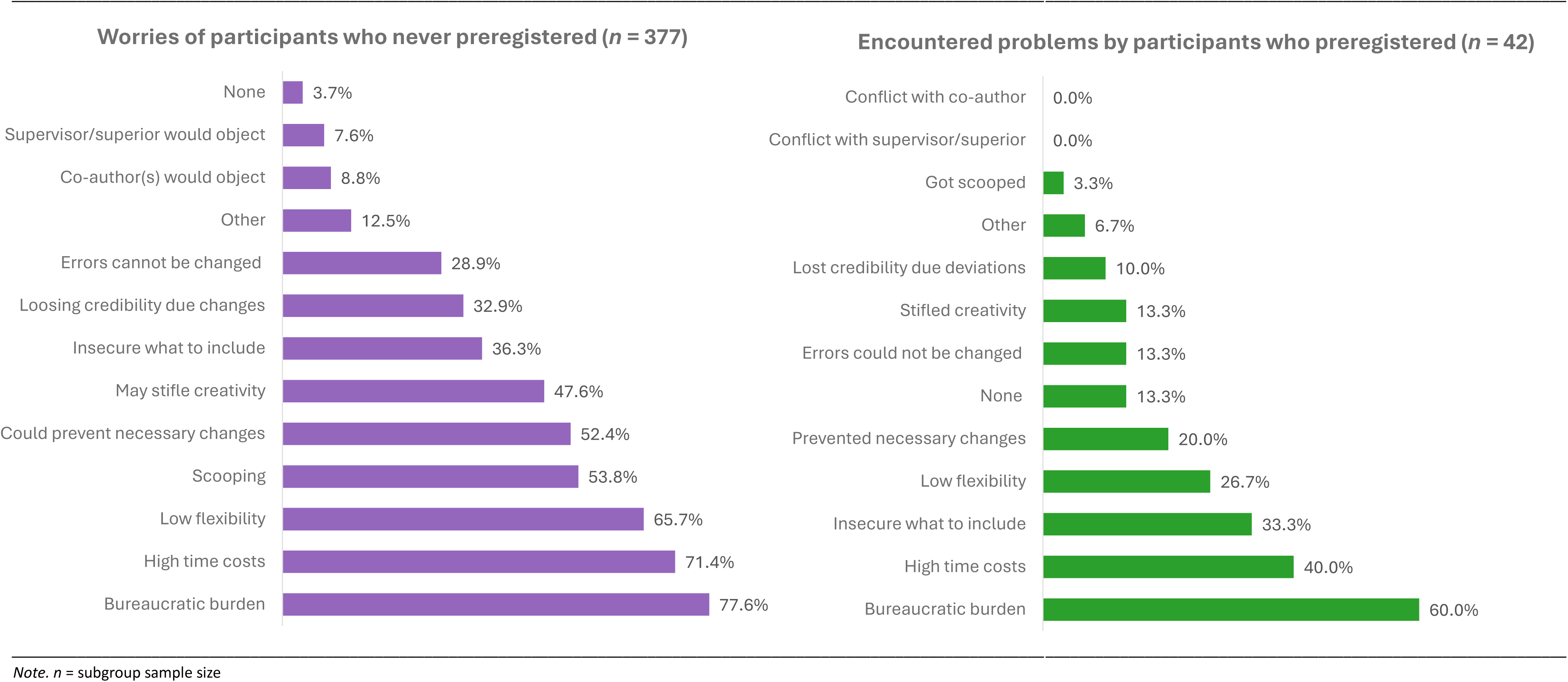
Barriers to Preregistration.

In addition to the closed-ended questions, open-ended responses on preregistration barriers were provided by up to 313 participants (S10 Table 1). The thematic analysis of these responses revealed four overarching themes: *practical and structural barriers*, *knowledge barriers*, *scientific barriers*, and *regulatory barriers* (Fig 2).

**Figure 2:**
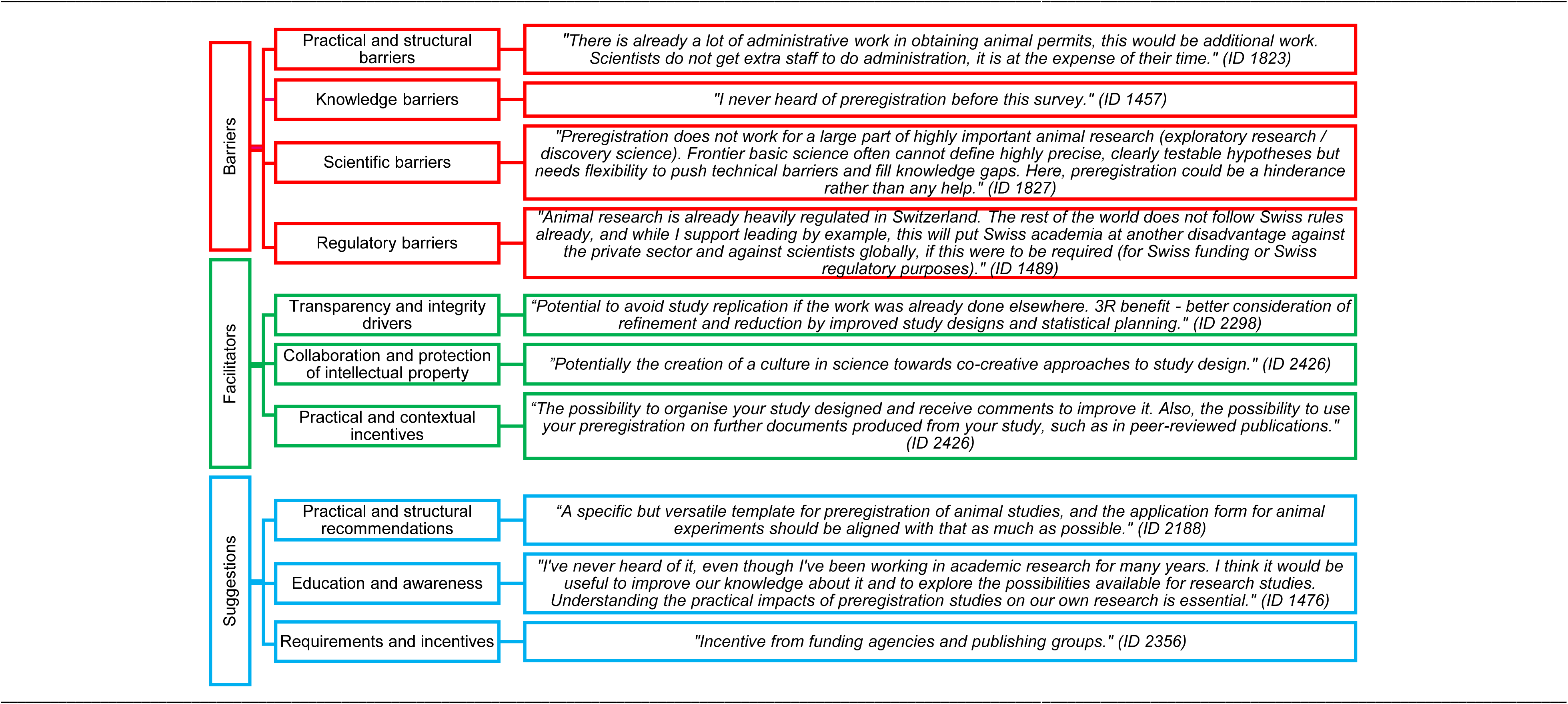
Thematic analysis.

##### Practical and structural barriers

Participants described both practical and structural challenges as barriers to preregistering their studies. Regarding practical barriers, they identified the overall bureaucratic burden, the time-consuming nature of the preregistration process, a lack of resources, and the increased costs and delays preregistration might entail. Concerns were also raised about the risk of scooping, confidentiality issues, and incompatibility with the industry practices and patent system, along with worries about external complaints and harassment. Structural barriers included the absence of institutional, journal, or funder requirements, as well as a lack of incentives or feedback mechanisms related to preregistration. In some cases, researchers reported limited support from co-authors and research partners for preregistering, while others were not in the position to make the decision themselves.

##### Knowledge barriers

Many participants stated that they were not aware that preregistration even existed. Some were unfamiliar with the procedure, describing a lack of guidance or uncertainty about the steps involved and the appropriate platforms for their studies. Others expressed general skepticism about the potential benefits and overall impact of preregistration in science.

##### Scientific barriers

Scientific barriers focused on concerns that preregistration stifles creativity, limits flexibility, and compromises innovation. Often, participants working in exploratory or fundamental research questioned the feasibility of preregistration for their studies. They emphasized the need to frequently deviate from their initial methodological plans as part of the iterative study designs necessary in their research, making preregistration difficult or unfeasible.

##### Regulatory barriers

A central issue under regulatory barriers was the overregulation of animal research in Switzerland. Some participants noted that the country’s highly controlled research environment limits the number of approved animal studies and reduces its international competitiveness. In some extreme cases, participants expressed concerns that additional practices, such as (mandatory) preregistration, could contribute to the relocation of Swiss research to countries with more permissive regulations.

#### Facilitators

When asked about the benefits of preregistration, up to 140 participants described various advantages of the practice (S10 Table 2). These perceived positive aspects could serve as motivators for adopting study preregistration in the future. Three main themes emerged from the thematic analysis: *transparency and integrity drivers*, *collaboration and protection of intellectual property*, and *practical and contextual incentives* (Fig 2). However, while some participants recognized these advantages, many saw no benefits at all, or found it appropriate only for confirmatory studies.

##### Transparency and integrity drivers

Despite the overall negative perception of preregistration, some participants pointed out that it could promote transparency in science, increase credibility and trustworthiness, and support the Open Science movement. Others highlighted its role in reducing parallel research and contributing to a better public understanding of the research world. In addition to transparency and openness, participants mentioned preregistration’s potential to promote scientific rigor and integrity. Some noted that it could help prevent questionable research practices, such as HARKing and p-hacking. Others explained that preregistration fosters thorough methodological planning and better study designs, while some described its role in mitigating publication bias, as all planned data collection is known in advance. Participants also suggested that preregistration may reduce researcher bias, enhance research quality, and promote less deviations and uniformity of experimental protocols. Furthermore, a few participants view preregistration as contributing to better consideration of animal use (3Rs) and improving the reliability and reproducibility of research findings.

##### Collaboration and protection of intellectual property

Participants also described collaboration-related benefits of preregistration. Some pointed out that preregistration may encourage scientific feedback and exchange, while others described its potential to foster collaborations and synergies. In addition, preregistration was seen by some as providing a clear track record of the studies, which could help protect researchers’ intellectual property.

##### Practical and contextual incentives

Finally, participants mentioned a few practical and contextual incentives related to preregistration that may facilitate its future use. Some noted that it may help with drafting papers and even speed up the journal review process, as parts of the manuscript are already written during the preregistration phase. Others highlighted the improved time and project management, which they attributed to better study planning required when preregistering research.

### Suggestions (Aim 5)

Up to 131 participants responded to the open-ended items regarding suggestions for improving preregistration (S10 Table 3). The following overarching themes emerged from the thematic analysis: *practical and structural recommendations*, *education and awareness recommendations*, and *incentive recommendations* (Fig 2).

#### Practical and structural recommendations

Participants offered a wide range of practical and structural recommendations to adapt the preregistration process to the Swiss context. They suggested the implementation of an easy-to-use tool with a clear, yet flexible, template and emphasized the need to make changes after preregistration. Some also proposed linking this tool to the license application procedure for animal studies and extending the embargo period to more than five years. In addition, participants asked for institutional support and resources, such as dedicated staff and funding, and highlighted the value of receiving expert feedback on the preregistered studies. Strong confidentiality measures were also recommended, and a few participants called for the international rather than national implementation of preregistration. Furthermore, many participants advocated for preregistration to remain voluntary in Switzerland, and if made mandatory, to apply it only to confirmatory studies.

#### Education and awareness recommendations

To facilitate the adoption of preregistration in Switzerland, participants described a need for more knowledge and awareness. Specifically, they requested information on where and how to preregister, concrete examples of preregistered studies, and access to workshops, courses, or continued education options. Furthermore, several participants emphasized the importance of proof that preregistration is beneficial across all types of research, including exploratory and fundamental studies.

#### Incentive recommendations

Principal investigators, institutions, funders, journals, and the scientific community were identified as essential in fostering the practice of preregistration. Several participants noted that endorsement or requirement from these parties would legitimize preregistration in animal research. Participants also listed a series of incentives that could encourage researchers to preregister, including faster licensing procedure for animal studies, benefits from funding organizations, institutional rewards, or publishing advantages.

## Discussion

This study represents the first attempt to evaluate the researchers’ experience with, and perspectives on, preregistration in animal research by examining the practices, attitudes and perceived challenges related to its implementation using the population of animal study directors accredited in Switzerland as a study population. To our knowledge, this is the first study worldwide evaluating psychosocial constructs related to preregistration among animal researchers.

### General experiences

We found that almost half of the participants were unaware of preregistration, and only 10% had preregistered a study before. These results are in strong contrast with a recent study conducted in the field of psychology in Germany (39), where only 4% of participants had never heard of preregistration, and 62% had preregistered at least one study in the past. Similarly, a further survey conducted in the Netherland reported that 43% of researchers across all research fields had prior experience with preregistration (44), while studies from behavioral and social sciences found rates of approximately 20% (45, 46). Although these differences may be partly explained by sampling biases inherent to the convenience samples of previous studies, our findings nonetheless suggest that preregistration awareness and experience are rather limited among animal researchers.

Interestingly, the motivations for first-time preregistration also differed across studies. In our sample, almost half of the participants indicated that they had preregistered because it was mandatory for a project, while only one third was self-motivated. By comparison, in the German study on psychologists, 49% had preregistered voluntarily, and only 13% reported preregistering because it was mandatory. This suggests that enforcement, rather than an intrinsic motivation, led Swiss animal scientists to preregister. In line with this, nearly half of the participants indicated that institutional policies could influence their decision to preregister in the future. However, almost a third stated that nothing could affect their decision. Our findings therefore suggest that policy changes may hold the greatest potential in promoting preregistration among animal researchers, but the widespread resistance within the research community should be considered when developing and implementing new policies.

### Psychosocial constructs and associations

On average, participants showed negative perceptions on all psychosocial constructs assessed here. They reported rather negative attitudes toward preregistration, perceived social norms as discouraging preregistration, and indicated limited abilities to preregister their own studies in terms of both resources and knowledge. Once again, our results differ from those described in other disciplines and countries, where preregistration was typically perceived and evaluated rather positively (39, 46, 47). Furthermore, both the intention and motivation to preregister future studies were low in our sample, while obstacles to preregister were perceived as relatively high. These findings indicate that animal researchers are reluctant to engage in preregistration and perceive the barriers to do so as substantial. However, participants with prior preregistration experience held more positive views of preregistration compared to those who had never preregistered a study, a pattern also observed in the field of psychology (39, 47). In contrast, participants who had worked more years in animal research expressed more negative perceptions of preregistration, aligning with evidence from behavioral science and psychology, where more experienced researchers showed greater reluctance toward preregistration (39, 46). Taken together, our findings underline substantial resistance and little support for preregistration within the Swiss animal research community, particularly among researchers who have never preregistered a study before and those with greater research seniority. Future efforts to promote preregistration should therefore tailor their approach to target these critical subgroups.

Interestingly, our analysis also revealed notable effects of gender and organization of employment on researchers’ perceived ability to preregister. Female researchers and participants employed in the private industry rated their knowledge required to preregister studies substantially lower than male researchers and those working in academic institutions. These findings may suggest potential structural differences in animal researchers’ capacities to preregister, with women and industry researchers perceiving themselves less equipped to engage in preregistration. However, the gender difference could also reflect that men tend to overestimate their knowledge more often than women, suggesting that women’s lower ratings may indicate more accurate or honest self-evaluations rather than a true lack of capacity.

There were also pronounced differences between those who preferred not to state their gender in the survey and those who did: this very small group of participants showed very negative attitudes, and very low intentions and motivations to preregister, but reported a much higher knowledge regarding preregistration. We can only speculate that these participants may have had negative experiences with preregistration and/or may have high privacy concerns that leaves them in general reluctant to disclose information, being it research ideas or private information such as their gender.

Lastly, the field of animal research in which participants were active also affected their perceptions of preregistration. Thus, researchers from General Biology (including such fields as animal behavior, nutrition, ecology and zoology) reported more favorable perceptions of and less competitive obstacles regarding preregistration than biomedical researchers. This might be due to a higher competitive pressure and resulting fear of scooping in these more applied fields of research. Such disciplinary differences should be considered when allocating resources toward fostering preregistration among all animal researchers.

### Barriers and facilitators

The most common practical and structural barriers identified by participants without preregistration experience included bureaucratic burden, excessive time investment, restricted flexibility, and challenges related to implementing necessary modifications to the study plan. However, the concerns anticipated by those without preregistration experience were substantially greater than the actual problems reported by participants who had preregistered before. These findings reveal a clear gap between perceived and actual barriers to preregistration, likely stemming from assumptions rather than practical realities of preregistration.

Targeted education for animal scientists may help address some of these perceived barriers as well as correct some misconceptions. Notably, aside from bureaucratic burden, similar barriers and discrepancies have been observed in psychology (39), revealing that these issues are shared across research fields. At the same time, it is paramount to acknowledge the existing administrative burden and the regulatory pressure faced by the animal research community in Switzerland. Before implementing new policies, authorities should be mindful of the potential resistance these changes may raise. Following the invitation to participate in the current study, the first author received several phone calls and emails from frustrated researchers expressing concerns about the possibility of an additional mandatory administrative requirement. Although these reactions were not always communicated diplomatically, they do reflect the general frustration and pressure experienced by Swiss animal scientists.

In addition to practical and structural barriers, several participants raised the issue of compatibility of preregistration with exploratory research. Differentiating between confirmatory and exploratory studies is essential to avoid promoting false-positive results as true effects (7, 46, 48). This delimitation could help reduce research waste by avoiding repetition and support the refinement of future studies, contributing to overall less animals use (24, 49). A common misunderstanding is that preregistration cannot be used in exploratory research and applies only to confirmatory studies. However, this barrier can be addressed by educating animal researchers about existing flexible preregistration templates (e.g., AsPredicted, OSF) that support the preregistration of their specific type of research.

Apart from the barriers to preregistration, participants also acknowledged a range of advantages. Although not always framed as direct facilitators, these perceived benefits could serve as motivators for adopting study preregistration. Participants saw value in preregistration for promoting transparency, improving study design and fostering scientific integrity. Some noted its potential to mitigate publication bias, enhance research quality and support better consideration of animal use. Emphasizing preregistration’s role in addressing questionable research practices and the overall contribution to scientific rigor might help convince researchers to engage more actively in study preregistration. Taken together, the barriers and facilitators observed in this study highlight the complexity of implementing preregistration in animal experimentation. Addressing misconceptions, emphasizing the benefits, and providing support for targeted problems may help reduce resistance and facilitate adoption of preregistration among animal researchers.

### Suggestions

Participants provided a wide range of recommendations for improving the preregistration process. Among practical aspects, they highlighted the need for a clear and flexible template, strong confidentiality measures, additional funding, specialized staff, and a connection between preregistration and the application procedure for animal experimentation. Moreover, participants stressed the importance of keeping preregistration voluntary. These suggestions underline the need to adapt preregistration to the setting-specific context (e.g. in terms of the authorization procedure) in order to enhance feasibility and promote broader acceptance among scientists. Given the limited knowledge about preregistration within the animal research community, participants also emphasized the need to increase awareness and offer education and training. Similar to findings in behavioral science (46), participants expressed uncertainty about the benefits of preregistration and highlighted the importance of evidence demonstrating that preregistration improves scientific quality. In addition, participants indicated a preference for concrete incentives to encourage preregistration, such as preregistration accelerating the authorization procedure, funding benefits, and publishing advantages. Providing the necessary infrastructure, appropriate educational options and a well-designed incentive system might thus help foster the use of preregistration in the animal research community.

### Strengths and limitations

A major strength of this study lies in the large response rate, with nearly one third of all accredited study directors of animal experiments in Switzerland participating in the survey. Moreover, study participants and non-responders appear similar with respect to the socio-demographic characteristics that were compared. Unlike previous studies, which were typically based on convenience samples and thus less representative of their target populations, our study reached the entire population of accredited study directors. Taken together, these aspects support the representativeness of the study sample and justify extrapolation of the findings to the overall population of animal scientists in Switzerland. Given that the Swiss animal research community is highly international and research cultures often vary more by field of research than by country, our findings may also offer valuable insights for the global animal research community. Particularly for countries with similar regulatory structures as those in Switzerland, the countries of the EU, our findings may provide useful guidance in view of fostering preregistration in animal research. Another important strength of the study is its mixed-methods design. The use of both closed-ended and open-ended measurements allowed for a deeper and more nuanced understanding of the studied constructs and enhances the robustness of the findings.

Apart from these strengths, a few limitations of the current study should be considered. First, despite the relatively large response rate, our sample might still have been affected by selection bias. Thus, individuals with particularly strong opinions about preregistration may have been more likely to participate in the study, leading to skewed outcomes. However, given that almost 40% of the participants had never heard of preregistration prior to the survey, it is rather unlikely that such strong views were held by many participants. Second, despite being introduced to the definition of preregistration at the beginning of the survey, some participants may not have read or fully understood the concept and may have confused it with the mandatory authorization procedure required to obtain a license for an animal experiment in Switzerland. Such misunderstanding would further reduce the proportion of participants with actual preregistration experience in our sample. This is supported by requests received after survey completion from three participants who asked to have their data deleted upon realizing that they had confused the two concepts. Finally, we do not know whether participants with preregistration experience had preregistered preclinical animal studies or clinical human studies, which may again lower the proportion of those with actual preregistration experience in animal research. As almost half of the participants indicated that their first preregistration was carried out because it was mandatory for a project, it is plausible that these cases may refer to clinical human studies where preregistration is legally required. Nevertheless, this may not compromise our conclusions, as the underlying concept and purpose of preregistration is the same across both research domains.

## Conclusion

Preregistration is arguably among the most impactful practices promoted by the Open Science movement. While it has long been established as standard practice in human clinical trials (17) and is increasingly taken up in other fields of research (45), preregistration has not been widely adopted in animal research. Given the limited knowledge of preregistration among animal scientists in Switzerland, as revealed by this study, comprehensive education and efforts to increase awareness are necessary before the research community would consider adopting preregistration. The widespread skepticism and resistance among specific subgroups of researchers – particularly those with no prior preregistration experience, greater research seniority, and those working in biomedical research – highlight the need for tailored strategies. Addressing key concerns, such as the bureaucratic burden and time constraints faced by animal researchers, and providing them with institutional support and adequate resources, would represent essential steps toward the successful implementation of preregistration in Switzerland and possibly elsewhere.

## Acknowledgments

We would like to acknowledge and thank Dr. Annamari Alitalo and Dr. med. vet. Otto Maissen for identifying the eligible study directors and carrying out the recruitment process for this study, as well as for their professional conduct and excellent communication in the administration of this research. We are also grateful to the AWO-Network for their support in issuing institutional reminders and facilitating participation. Finally, we would like to thank all participants who took the time to complete our survey.

