## Supplemental material for "Researchers‘ perspectives on preregistration in animal research"

### S1 Table: Deviations from preregistration plan

| Deviations | | | | | |
|---|---|---|---|---|---|
| | Details | | Original Wording | Deviation Description | Reader Impact |
| 1 | Type | Analysis | Participants who will not respond correctly to the attention check will be excluded from the analysis (*This is an attention check question. Please select both "Agree" and "Strongly agree" for this item to demonstrate that you are paying attention.).* Additionally, participants who complete the survey more than once using the same entry code will be excluded from the analysis, except for their first completed attempt. | Participants with more than 90% missing data were also removed from the analysis. | The deviation has no impact on the study findings, as the incomplete responses were unusable for statistical analysis and would not have contributed meaningfully to the results. |
| | Reason | New knowledge | | | |
| | Timing | After data access | | | |
| 2 | Type | Variables | The items used in the survey are adapted from the questionnaire developed by Spitzer and Mueller, who report high reliabilities for all six scales: attitudes towards study preregistration, subjective norms, perceived behavioral control, intentions, motivations, and obstacles with regard to study preregistration. However, it is not clear whether these scales are really unidimensional. Therefore, we will conduct Principal | The PCA analysis revealed that two of the six original scales exhibited a two-component structure. Consequently, each was divided into two subscales:<br>• Perceived Behavioral Control Scale:<br> ○ Resources Subscale<br> ○ Knowledge Subscale<br>• Obstacles Scale<br> ○ Practical Obstacles Subscale<br> ○ Competitive Obstacles Subscale | The use of the two new subscales improved the psychometric quality of the instrument and allowed for a closer conceptual alignment with the tested constructs. This refinement increased the precision of the questionnaire and led to a more robust analysis. |
| | Reason | New knowledge | | | |
| | Timing | After data access | | | |

| | | | | | |
|---|---|---|---|---|---|
| | | | Component Analyses (PCA) to explore the underlying structure of the items in the survey within each of the six outcomes. To arrive at an appropriate number of components we will use the scree criterion together with the parallel analysis criterion (using the eigenvalue means of 1000 random samples for comparison). We generally expect the scales to be unidimensional, but in case that there is more than one component according to the above criteria, the extraction will be followed by an oblique Oblimin-rotation to arrive at a simple structure even in case of correlated components. | | |
| 3 | Type | Analysis | For the association analysis, we will conduct simple and multiple linear regression models using the scale scores for attitudes, subjective norms, perceived behavioral control, intentions, motivations, and obstacles as dependent variables. To explore bivariate associations between the predictors and the dependent variables, we will run simple regressions (one predictor at a time) for each of the six | Simple and multiple linear regression models with the Preregistration Scale scores as dependent variables were initially considered for the association analysis and preregistered on OSF. However, given that three categorical predictors (gender, field of animal research, organization of employment) included more than two levels, the analysis was adapted. Instead of multiple linear regressions, a | This new approach allowed for the detection of specific group differences on each of the Preregistration Scale, which would not have been possible using multiple linear regression models. We therefore were able to see differences between all groups of the predictors:<br>• Gender (“female” vs. “male” vs. “prefer not to say”)<br>• Field of animal research (“basic and experimental medical research” vs. |

| | | | | | |
|---|---|---|---|---|---|
| | | | dependent variables separately. The predictors included in the univariable linear regression models will be:<br>• Gender ("male" vs. "female")<br>• Research experience ("animal research experience" in years as continuous variable)<br>• Preregistration experience ("prior experience with preregistration" vs. "no experience with preregistration")<br>• Field of animal research ("basic biology research" vs. "general biology" vs. "basic and experimental research")<br>• Type of institution ("academia vs. non-academia").<br>In a subsequent step, we will conduct six separate multiple linear regression models (one for each dependent variable) and we will fit all five predictors in the regression models independent of their significance in the univariable analysis. This is done to explore the unique associations of each predictor with the dependent variables. Unstandardized and standardized partial regression coefficients as well as the squared semipartial correlations ($\Delta R^2$) will be reported. | multivariate analysis of covariance (MANCOVA) was conducted, followed by univariate analyses of variance (ANCOVAs) and post hoc comparisons. | "general biology" vs. "basic biology research")<br>• Organization of employment ("academic" vs. "private" vs. "other") |

**Unregistered Steps**

| | Details | | Original Wording | Unregistered Step Description | Reader Impact |
|---|---|---|---|---|---|
| 1 | Type | Analysis | - | An unplanned comparison of sociodemographic characteristics was conducted between participants who had preregistered and those who never preregistered a study, due to the large sample size difference between the groups. Two-sample t-tests were used for mean differences and Fisher's Exact Test for differences in categorical variables. | This unplanned analysis just gives more context about the sociodemographic differences between the two groups and has no impact on the readers interpretation of the study results |
| | Timing | After data access | | | |

### S3 Image: Attention check item

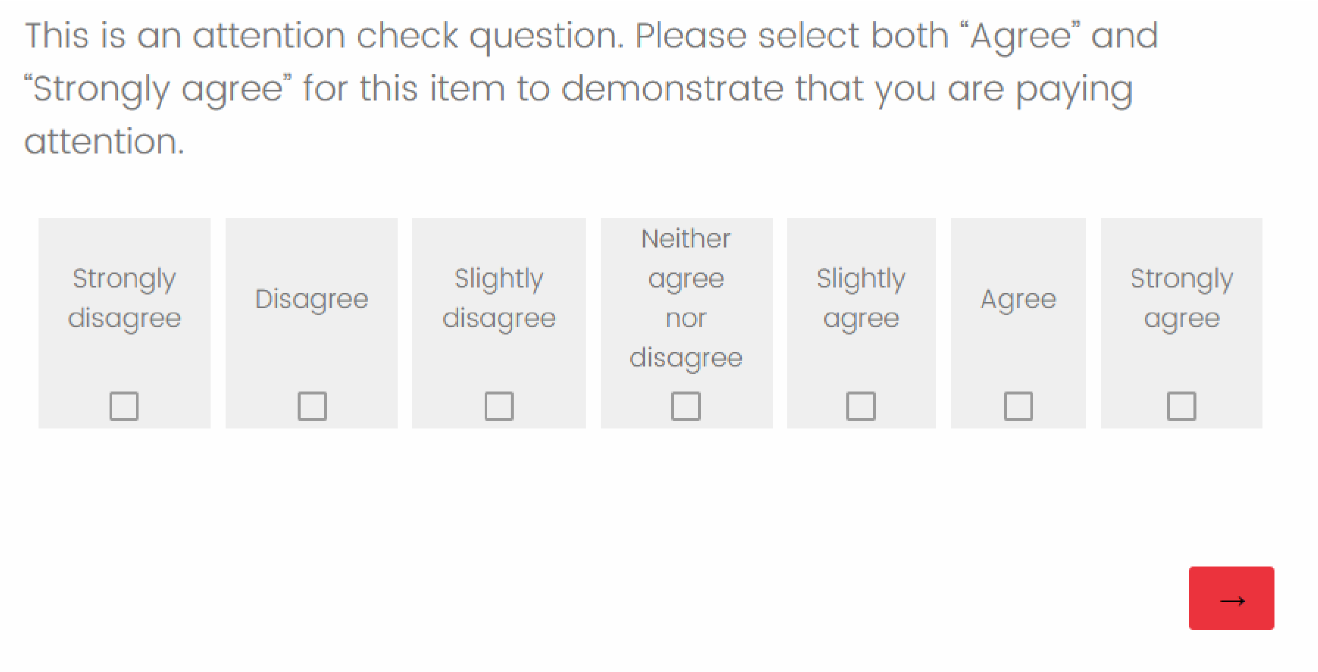

This is an attention check question. Please select both “Agree” and “Strongly agree” for this item to demonstrate that you are paying attention.

| Strongly disagree | Disagree | Slightly disagree | Neither agree nor disagree | Slightly agree | Agree | Strongly agree |
|---|---|---|---|---|---|---|
| ☐ | ☐ | ☐ | ☐ | ☐ | ☐ | ☐ |

→

### S4 Table: General experiences with study preregistration

| **ALL PARTICIPANTS (*N* = 418)** | **% (*n*)** |
|---|---|
| ***Have you ever preregistered a study?*** | |
| Yes | 10.0% (42) |
| No | 90.0% (376) |
| *N* | 418 (0 missing) |
| ***Where did you learn about preregistration?*** | |
| Never heard before this survey | 39.2% (147) |
| Read about it | 31.5% (118) |
| Further education | 27.2% (102) |
| Conversation with colleagues | 19.7% (74) |
| Presentation at conference(s) | 12.0% (45) |
| During my studies | 5.9% (22) |
| Information event at workplace | 5.3% (20) |
| Do not remember | 5.3% (20) |
| Other | 2.1% (8) |
| Supervisor/superior | 1.6% (6) |
| Co-author(s) | 1.1% (4) |
| *N* | 374 (44 missing) |
| ***Which persons or institutions have an influence in your decision to preregister or not to preregister your study?*** | |
| Institutional policies | 47.6% (175) |
| Funding guidelines | 35.6% (131) |
| Supervisor/superior | 32.6% (120) |
| Nobody | 28.0% (103) |
| Journal guidelines | 26.4% (97) |
| Co-author(s) | 19.0% (70) |
| Colleagues | 15.8% (58) |
| Guidelines of learned societies | 9.0% (33) |
| Other | 1.6% (6) |
| *N* | 360 (58 missing) |
| **ONLY IF PARTICIPANTS PREREGISTERED BEFORE (n = 42)** | **% (*n*)** |
| ***Approximately how many studies have you preregistered before?*** | |
| *M (SD)* | 7.6 (13.62) |
| *Mdn* | 3.0 |
| Range | 1 - 80 |
| *n* | 38 (4 missing) |
| ***On average, how many hours would it take you to preregister a study?*** | |
| *M (SD)* | 22.9 (32.66) |
| *Mdn* | 11.0 |
| Range | 1 - 150 |
| *n* | 38 (4 missing) |
| ***What was/were the reason/s for your first preregistration? Please select every option that applies.*** | |
| Mandatory for a project | 45.5 % (15) |
| Self-motivated | 27.3 % (9) |
| Suggestion from supervisor/superior | 21.2 % (7) |

| | |
|---|---|
| Recommendation by co-authors | 18.2 % (6) |
| Requirement to get funding | 15.2 % (5) |
| Conversation with colleagues | 6.1 % (2) |
| Other | 3.0 % (1) |
| *n* | 33 (9 missing) |
| ***When I preregister, I use the following repository (i.e., uploading platform) for sharing my preregistration. Please select every option that applies.*** | |
| animalstudyregistry.org (ASR) | 29.6% (8) |
| Offline (e.g., I only share it with my co-authors or store it at my institution) | 22.2% (6) |
| ClinicalTrials.gov | 18.5% (5) |
| Institutional platform | 14.8% (4) |
| osf.io (Open Science Framework) | 11.1% (3) |
| Other | 7.4% (2) |
| AsPredicted.org | 7.4% (2) |
| PROSPERO | 3.7% (1) |
| Registered reports with a specific journal | 3.7% (1) |
| preclinicaltrials.eu | 0.0% (0) |
| researchregistry.com | 0.0% (0) |
| Personal website | 0.0% (0) |
| *n* | 27 (15 missing) |
| ***What is your preferred preregistration template(s) (i.e., form that lists important elements to preregister and can be used by researchers to create their own preregistration)?*** | |
| No preference | 52.0% (13) |
| animalstudyregistry.org (ASR) | 20.0% (5) |
| Other | 8.0% (2) |
| ClinicalTrials.gov | 8.0% (2) |
| PREPARE checklist | 8.0% (2) |
| Do not use templates | 4.0% (1) |
| osf.io (Open Science Framework) | 4.0% (1) |
| preclinicaltrials.eu | 4.0% (1) |
| AsPredicted.org | 0.0% (0) |
| PROSPERO | 0.0% (0) |
| researchregistry.com | 0.0% (0) |
| *n* | 25 (17 missing) |
| ***Why do you prefer the selected template(s)?*** | |
| Comprehensive | 50.0% (6) |
| Fits well with my research area | 50.0% (6) |
| Easy to use | 41.7% (5) |
| Other | 25.0% (3) |
| Time-efficient | 25.0% (3) |
| Preferred by colleagues | 16.7% (2) |
| The only template I know | 16.7% (2) |
| First template I used | 16.7% (2) |
| Recommended by co-author(s) | 8.3% (1) |

| | |
|---|---|
| Recommended by supervisor/superior | 0.0% (0) |
| *n* | 30 (12 missing) |

*Note. M* = mean; *SD* = standard deviation; *Mdn* = median; *N* = total sample size; *n* = subgroup sample size.

### S5 Table: Subsample comparison – Preregistration experience vs. no preregistration experience

| Characteristics | Preregistration experience (*n* = 42) | No preregistration experience (*n* = 376) | p |
|---|---|---|---|
| **Age**[a] | | | |
| *M (SD)* | 45.9 (10.49) | 47.0 (9.05) | .496 |
| *Mdn* | 42.0 | 46.0 | |
| Range | 31 - 74 | 26 - 65 | |
| *n* | 41 (1 missing) | 362 (14 missing) | |
| **Gender** | | | |
| Female | 42.9% (18) | 40.8% (152) | .758 |
| Male | 52.4% (22) | 55.8% (208) | |
| Other / Prefer not to say | 4.8% (2) | 3.5% (13) | |
| *n* | 42 (0 missing) | 373 (3 missing) | |
| **Years registered as study director**[a] | | | |
| *M (SD)* | 10.2 (8.34) | 10.1 (7.90) | .938 |
| *Mdn* | 7.0 | 8.0 | |
| Range | 0 - 30 | 0 - 42 | |
| *n* | 41 (1 missing) | 363 (13 missing) | |
| **Years in animal research** | | | |
| *M (SD)* | 19.4 (8.79) | 20.2 (8.85) | .605 |
| *Mdn* | 17.5 | 19.0 | |
| Range | 7 - 36 | 2 - 44 | |
| *n* | 42 (0 missing) | 372 (4 missing) | |
| **Academic age** | | | |
| *M (SD)* | 15.7 (9.95) | 17.5 (9.24) | .292 |
| *Mdn* | 12.0 | 16.5 | |
| Range | 1 - 37 | 0 - 40 | |
| *n* | 42 (0 missing) | 366 (10 missing) | |
| **Educational attainment** | | | |
| Master's degree | 2.4% (1) | 5.6% (21) | .645 |
| PhD / Dr. med. | 52.4% (22) | 51.5% (192) | |
| Habilitation / professorship | 42.9% (18) | 41.8% (156) | |
| Other | 2.4% (1) | 1.1% (4) | |
| *n* | 42 (0 missing) | 373 (3 missing) | |
| **Seniority level** | | | |
| PhD student / technician | 0.0% (0) | 4.3% (16) | .282 |
| Postdoc. / senior researcher | 33.3% (14) | 25.9% (96) | |
| Lecturer | 2.4% (1) | 2.7% (10) | |
| Group leader / Professor | 52.4% (22) | 60.9% (226) | |
| Other | 11.9% (5) | 6.2% (23) | |
| *n* | 42 (0 missing) | 372 (5 missing) | |
| **Field of animal research** | | | |
| Basic biological research[b] | 4.8% (2) | 14.4% (54) | .140 |
| General biology[c] | 19.0% (8) | 13.4% (50) | |

| | | | |
|---|---|---|---|
| Basic and experimental medical research[d] | 76.2% (32) | 72.2% (270) | |
| *n* | 42 (0 missing) | 374 (2 missing) | |
| **Organization of employment** | | | |
| Academic | 69.0% (29) | 78.6% (294) | .119 |
| Academic & other | 0.0% (0) | 2.4% (9) | |
| Private | 23.8% (10) | 12.6% (47) | |
| Governmental | 7.1% (3) | 2.1% (8) | |
| Non-profit | 0.0% (0) | 4.3% (16) | |
| *n* | 42 (0 missing) | 374 (2 missing) | |

*Note. M* = mean; *SD* = standard deviation; *Mdn* = median; *n* = subgroup sample size.

[a] In contrast to Table 1 in the main manuscript, the values displayed here are based on our own data collection and not on the data provided by the Swiss Federal Food Safety and Veterinary Office (FSVO).

[b] Including following fields: Biomedical Engineering; Cancer Research; Cardiovascular Research; Endocrinology; Immunology; Medical Microbiology; Neuroscience; Nutrition and Metabolism; Pathology and/or Pathophysiology; Pharmacology; Physiology; Regenerative Medicine; Toxicology; Veterinary Medicine; Virology.

[c] Including following fields: Animal Breeding; Animal Nutrition; Animal Welfare; Ecology; Ethology; Evolution; Laboratory Animal Science; Wildlife Biology; Zoology.

[d] Including following fields: Biochemistry; Biophysics; Cell Biology; Cytology; Developmental Biology; Embryology; Epigenetics; Experimental Microbiology; Genetics; Molecular Biology; Radiobiology; Structural Biology.

Significant results are marked with *p <0.05, **p <0.01, ***p <0.001.

### S6 Psychometric quality

All psychosocial construct scales were assumed to be unidimensional, but this assumption has not been examined in prior research. We therefore tried to establish strict or essential unidimensionality of the attitudes, perceived social norms, perceived behavioral control, intentions, motivations, and obstacles towards preregistration scales. While strict unidimensionality refers to the situation where there truly exists only one underlying dimension for a scale or construct, essential unidimensionality refers to the situation where there is a dominant general factor or component but nevertheless there is some sort of multidimensionality in terms of subdimensions present. It is important to establish whether the general factor is strong enough to allow for using an overall composite scale score for the construct.

Several rules and indices have been suggested to assess essential unidimensionality. Slocum-Gori and Zumbo (1) recommend a combination of parallel analysis and a criterion based on the ratio of the first to the second eigenvalue: if this ration is greater than three or (in a stricter version) greater than four - thus, when the first eigenvalue is at least three times as high as the second eigenvalue, essential unidimensionality can be assumed even when parallel analysis suggests the existence of more than one component.

In addition, following McDonald (2) and Revelle and Zinbarg (3), we computed omega hierarchical ($\omega_H$) from a bifactor model to assess essential unidimensionality. A bifactor structure means that there is a general factor indicating unidimensionality, but also two or more so-called group factors that are modelled simultaneously. While Cronbach's alpha and also the "normal" coefficient ω represent the proportion of shared variance in the data vis-à-vis to total variance, $\omega_H$ represents the proportion of variance in overall scores that can be attributed to a single general factor, while variation in scores coming from the existence of group factors is treated as measurement error (4). Values of $\omega_H$ above .70 are generally taken as evidence that a scale is sufficiently unidimensional for practical purposes.

For scales that showed clear (strict) unidimensionality (we assumed this when parallel analysis suggested only one factor) we used coefficient ω (and not α) as a measure of reliability. The reason is that alpha's assumption of an essential tau-equivalent model (implying equal factor loadings for all items) is seldom warranted and can lead to an underestimation of reliability when loadings are not equal for all items (congeneric model). On the other hand, ω is suitable for this case since it is computed directly from the factor loadings of a unidimensional solution.

### Attitudes Scale

A parallel analysis (based on principal components analysis, from now abbreviated as PA-PCA) indicated a two-component structure, accounting for 58.5% of the total variance (Component 1 = 51.8%, Component 2 = 6.7%) of attitudes towards preregistration. The eigenvalues were 11.92 and 1.54 for the first and second component, respectively, with a ratio of the first to the second eigenvalue of 7.72. The eigenvalue of the second component was only slightly above chance level according to the PA-PCA. Coefficient $\omega_H$ was .95. These two criteria strongly suggest essential unidimensionality of the Attitudes Scale.

A one-component PCA was computed (see Table 1) where most items loaded strongly (>.70) on the single component. However, Items 9 and 21 showed low factor loadings (.24 and .14) as well as low corrected item-total correlations (.22 and .13). In addition, upon closer examination, both items did not align well conceptually with the construct of attitudes and were therefore excluded to obtain a highly homogenous Attitudes Scale. The final revised 21-item Attitudes Scale showed excellent reliability (ω = .96) and was used in all subsequent analyses reported in the main manuscript.

**Table 1: Psychometric properties and component loadings of the Attitude Scale (23 items)**

| Items | *N* | *M* | *SD* | r.drop[a] | Component Loading |
|---|---|---|---|---|---|
| 1. Preregistration is important to me. | 370 | -1.34 | 1.61 | 0.79 | 0.83 |
| 2. I have more trust in research findings when the study has been preregistered. | 369 | -1.18 | 1.68 | 0.81 | 0.84 |
| 3. I have more trust in researchers who preregister their studies than in those who do not. | 371 | -1.26 | 1.67 | 0.81 | 0.84 |
| 4. My field of science benefits from preregistration. | 370 | -1.12 | 1.67 | 0.81 | 0.84 |
| 5. Preregistration should be mandatory for all types of studies. | 370 | -1.80 | 1.45 | 0.67 | 0.71 |
| 6. Preregistration should be mandatory only for specific types of studies (e.g. confirmatory studies). | 368 | -0.25 | 1.85 | 0.47 | 0.51 |
| 7. Preregistration hinders exploratory/discovery research. (R)[b] | 367 | -1.32 | 1.62 | 0.51 | 0.55 |
| 8. Preregistration should be an evaluation criterion in selection decisions (e.g. publication, research evaluation, funding etc.). | 368 | -1.50 | 1.61 | 0.76 | 0.79 |
| 9. Preregistration can be exploited (e.g. by cheating). (R)[b] | 365 | -0.70 | 1.37 | 0.22 | 0.24 |
| 10. Preregistration does not improve research substantially. (R)[b] | 368 | -1.02 | 1.60 | 0.80 | 0.82 |
| 11. Preregistration cannot prevent questionable research practices. (R)[b] | 364 | -1.42 | 1.46 | 0.55 | 0.59 |
| 12. Preregistration can prevent selective reporting (i.e., only reporting significant | 364 | -0.03 | 1.78 | 0.56 | 0.59 |

| | | | | | |
|---|---|---|---|---|---|
| results or results compatible with the hypotheses). | | | | | |
| 13. Preregistration can prevent p-hacking (i.e., misusing data analyses to find patterns that can be presented as statistically significant). | 360 | -0.37 | 1.64 | 0.64 | 0.66 |
| 14. Preregistration can prevent HARKing (i.e., adjusting hypotheses to the observed results). | 360 | -0.03 | 1.74 | 0.69 | 0.71 |
| 15. Preregistration can prevent publication bias (i.e., only publishing studies with positive/significant results). | 360 | -0.49 | 1.83 | 0.70 | 0.73 |
| 16. Preregistration improves a study's quality. | 360 | -0.78 | 1.68 | 0.82 | 0.85 |
| 17. Preregistration increases the credibility of animal research. | 358 | -0.72 | 1.76 | 0.81 | 0.83 |
| 18. The costs of preregistering a study are higher than the benefits. (R)[b] | 355 | -1.09 | 1.55 | 0.63 | 0.67 |
| 19. Preregistering studies is generally unnecessary. (R)[b] | 355 | -0.71 | 1.70 | 0.76 | 0.79 |
| 20. Preregistration makes science more transparent. | 356 | -0.02 | 1.79 | 0.79 | 0.82 |
| 21. There is a large variety of tools available to create preregistrations (for example templates and repositories). | 329 | 0.03 | 0.72 | 0.13 | 0.14 |
| 22. Preregistration is not useful in practice. (R)[b] | 350 | -0.92 | 1.54 | 0.79 | 0.82 |
| 23. A preregistration badge (i.e., a public acknowledgment that a study was preregistered provided by many journals) increases my trust in a study. | 352 | -0.81 | 1.68 | 0.79 | 0.81 |

*Note.* *N* = number of participants who answered the item; *M* = mean; *SD* = standard deviation.

[a]Corrected item-total correlation, indicating how well each item correlates with the total scale score when that item is excluded.

[b]Reverse-coded item.

### Subjective Norms Scale

The PA-PCA indicated a one-component structure, accounting for 48.9% of the total variance of subjective norms regarding preregistration. The ratio of the first to the second eigenvalue was 3.15 (with eigenvalues of the first two components of 3.42 and 1.09, respectively). Coefficient $\omega_H$ was .80. Both criteria suggest unidimensionality of the Subjective Norms Scale.

A one-component PCA was computed (Table 2). All but two items loaded strongly (>.70) on the component: Item 6 had a loading of .60., while Item 2 showed a very low component loading of .13. Moreover, Item 2 did not conceptually align with the construct of subjective norms and was therefore excluded to obtain a more homogenous scale. The final revised 6-item Subjective Norms Scale showed a good reliability of ω = .84 and was further used in the analyses reported in the main paper.

**Table 2: Psychometric properties and component loadings of the Subjective Norms Scale (7 items)**

| Items | *N* | *M* | *SD* | r.drop[a] | Component Loading |
|---|---|---|---|---|---|
| 1. My peers or colleagues motivate me to preregister my studies. | 346 | -1.58 | 1.40 | 0.67 | 0.79 |
| 2. I want to be part of the Open Science Movement. | 346 | 0.94 | 1.54 | 0.10 | 0.13 |
| 3. My supervisor/superior wants me to preregister my studies. | 331 | -1.14 | 1.36 | 0.60 | 0.73 |
| 4. My co-authors want me to preregister my studies. | 341 | -1.33 | 1.36 | 0.71 | 0.84 |
| 5. Preregistration is highly acknowledged in my research community. | 346 | -1.66 | 1.35 | 0.69 | 0.83 |
| 6. I feel social pressure to preregister my studies. | 349 | -1.47 | 1.47 | 0.45 | 0.60 |
| 7. I think that most researchers in my field preregister their studies. | 349 | -2.15 | 1.10 | 0.52 | 0.70 |

*Note. N* = number of participants who answered the item; *M* = mean; *SD* = standard deviation.

[a]Corrected item-total correlation, indicating how well each item correlates with the total scale score when that item is excluded.

### Perceived Behavioral Control Scale

The PA-PCA suggested a three-component solution. The three components accounted for 74.2% of the total variance, with eigenvalues of 2.28, 1.73, and 1.18 for Components 1 through 3, respectively. The ratio of the first to the second eigenvalue was 1.32, and the coefficient $\omega_H$ was .46. Thus, unidimensionality could not be assumed for the Perceived Behavioral Control Scale.

Since the third eigenvalue (1.18) was only slightly above chance level in the PA-PCA, and because the loading pattern of the three-component oblimin-rotated solution was inconclusive with two items showing substantial cross-loadings, a two-component solution was considered. This solution explained 57.4% of the overall variance and showed a clear simple structure (Table 3). The first component, with high loadings of items 1, 4, 5, and 6, represented the availability of resources and support from colleagues and supervisors for preregistration and was labeled *Perceived Behavioral Control – Resources Subscale*. The second component, with high loadings of items 2, 3, and 7, represented the knowledge needed for preregistration (two items) and the authority to decide about preregistration (one item), and was named *Perceived Behavioral Control – Knowledge Subscale*. While Perceived Behavioral Control – Resources Subscale (4 items) showed acceptable reliability with ω = .72, the reliability estimate of the Perceived Behavioral Control – Knowledge Subscale (3 items) was questionable with ω = .65 but that we accepted for our exploratory study. Thus, both subscales were ultimately used in the analyses presented in the main manuscript.

**Table 3: Psychometric properties and component loadings of the Perceived Behavioral Control Scale (7 items)**

| Items | *N* | *M* | *SD* | Component Loading 1 | Component Loading 2 |
|---|---|---|---|---|---|
| 1. It is or it would be easy for me to preregister my studies. | 348 | -1.3 | 1.4 | 0.68 | -0.18 |
| 2. I know how to create and upload a preregistration. | 346 | -1.3 | 1.6 | 0.17 | 0.71 |
| 3. It is my decision to preregister my studies or not. | 351 | 0.88 | 1.8 | -0.12 | 0.65 |
| 4. I do not have the time nor the resources to preregister my studies. (R)[a] | 347 | -1.19 | 1.5 | 0.70 | -0.19 |
| 5. My supervisor/superior does not support preregistration. (R)[a] | 330 | -0.12 | 1.4 | 0.78 | 0.25 |
| 6. My co-authors do not support preregistration. (R)[a] | 336 | -0.22 | 1.3 | 0.82 | 0.02 |
| 7. I do not feel well informed about preregistration. (R)[a] | 347 | -0.84 | 1.7 | -0.05 | 0.83 |

*Note. N* = number of participants who answered the item; *M* = mean; *SD* = standard deviation.

[a]Reverse-coded item.

### Intentions Scale

The PA-PCA of the 3-item Intentions Scale indicated a single underlying component with an eigenvalue of 2.46. The one component explained 82% of the total variance with high factor loadings for all three items. Coefficient $\omega_H$ was .89. These values support the appropriateness of the one-component solution, and all three items were used in the calculation of the scale score. The reliability estimate was $\omega$ = .89.

**Table 4: Psychometric properties and component loadings of the Intention Scale (3 items)**

| Items | *N* | *M* | *SD* | r.drop[a] | Component Loading |
|---|---|---|---|---|---|
| 1. I will preregister my studies in the future. | 346 | -1.16 | 1.57 | 0.81 | 0.92 |
| 2. I am open to the idea of preregistering my research in the future. | 348 | -0.43 | 1.96 | 0.81 | 0.92 |
| 3. I do not intend to preregister my future study/studies. (R)[b] | 347 | -0.69 | 1.91 | 0.73 | 0.87 |

*Note. N* = number of participants who answered the item; *M* = mean; *SD* = standard deviation.

[a]Corrected item-total correlation, indicating how well each item correlates with the total scale score when that item is excluded.

[b]Reverse-coded item.

### Motivations Scale

For the 10-items Motivations Scale, a single-component structure was suggested by the PA-PCA. The eigenvalue of the single component was 6.54 and it thus represented 65.4% of the total variance. Factor loadings were high for most items (only two items loaded below .80, see Table 5), supporting the unidimensionality of the Motivations Scale.

The unidimensionality was confirmed by a high omega hierarchical ($\omega_H$ = .94) and the reliability estimated by coefficient omega was also excellent ($\omega$ = .94).

**Table 5: Psychometric properties and component loadings of the Motivations Scale (10 items)**

| Items | *N* | *M* | *SD* | r.drop[a] | Component Loading |
|---|---|---|---|---|---|
| 1. I feel like preregistration is an investment in my future (e.g., it is helpful for my credibility or career). | 347 | -1.29 | 1.56 | 0.81 | 0.86 |
| 2. I believe that it is becoming harder to publish studies that were not preregistered. | 346 | -1.24 | 1.50 | 0.50 | 0.57 |
| 3. Preregistration helps plan studies better. | 348 | -0.65 | 1.84 | 0.77 | 0.82 |
| 4. The preregistration badge (i.e., a public acknowledgment that a study was preregistered provided by many journals) would be an incentive for me to preregister my studies. | 348 | -0.76 | 1.68 | 0.76 | 0.81 |
| 5. I feel morally compelled to preregister my studies. | 347 | -1.36 | 1.51 | 0.76 | 0.81 |
| 6. Preregistration makes studies more transparent. | 348 | -0.46 | 1.86 | 0.81 | 0.85 |
| 7. Preregistration makes studies more trustworthy. | 347 | -0.93 | 1.72 | 0.87 | 0.90 |
| 8. I want others to be able to comment on my planned studies. | 347 | -0.73 | 1.72 | 0.62 | 0.68 |
| 9. Preregistration helps researchers protect themselves from their own biases. | 346 | -0.45 | 1.71 | 0.82 | 0.86 |
| 10. Preregistration represents good scientific practice. | 347 | -0.38 | 1.68 | 0.82 | 0.87 |

*Note. N* = number of participants who answered the item; *M* = mean; *SD* = standard deviation.

[a]Corrected item-total correlation, indicating how well each item correlates with the total scale score when that item is excluded.

#### Obstacles Scale

A PA-PCA on the 10-item Obstacle Scale revealed a two-component structure (eigenvalues of 3.79 and 1.60), explaining 53.9% of the total variance. Although the ratio of first to the second eigenvalue (3.79/1.60 = 2.37) was below the cutoff of 3 for essential unidimensionality, this assumption was supported by omega hierarchical ($\omega_H$ = .73).

Since the evidence was not clearly in favor of essential unidimensionality, we adopted the two-component solution presented in Table 6. Items 3 to 6 and 9 and 10 had high loadings on the first component. This dimension reflected practical obstacles to preregistration, particularly disadvantages in terms of time and effort, and was labeled *Practical Obstacles Subscale*. The second component with substantial to high loadings of items 1, 2, 7 and 8 represented concerns about scooping, confidentiality, and competitive disadvantages, and was labeled *Competitive Obstacles Subscale*.

The Practical Obstacles Subscale (6 items) showed a good reliability (ω = .84), whereas the Competitive Obstacles Subscale (4 items) demonstrated a reliability that was questionable (ω = .66) but that we deemed acceptable for our exploratory purposes. For the analyses reported in the main manuscript, we used both newly developed subscales.

**Table 6: Psychometric properties and component loadings of the Obstacle Scale (10 items)**

| Items | N | *M* | *SD* | Component Loading 1 | Component Loading 2 |
|---|---|---|---|---|---|
| 1. Preregistration puts me at a disadvantage in comparison to those who do not preregister. | 346 | 0.29 | 1.6 | 0.23 | 0.52 |
| 2. I am concerned that after pre-registering my studies others will find errors in, or deviations from, my study plans. | 346 | -0.37 | 1.5 | -0.11 | 0.43 |
| 3. I do not like that preregistration limits my flexibility in research. | 346 | 1.43 | 1.5 | 0.62 | 0.20 |
| 4. Preregistration incurs a considerable time cost. | 345 | 1.82 | 1.1 | 0.84 | -0.06 |
| 5. Preregistration is a bureaucratic exercise. | 347 | 1.67 | 1.4 | 0.81 | 0.08 |
| 6. For me, there are not enough incentives to preregister my studies. | 346 | 1.33 | 1.4 | 0.50 | 0.10 |
| 7. I would be afraid of scooping (i.e., someone taking my idea and publishing it before me) when preregistering my study. | 348 | 1.20 | 1.6 | -0.01 | 0.85 |
| 8. I would be unsure about confidentiality issues and intellectual property rights when preregistering. | 347 | 1.50 | 1.5 | -0.01 | 0.83 |
| 9. For my projects, preregistration is unnecessary. | 345 | 1.35 | 1.6 | 0.81 | -0.12 |
| 10. Preregistration would slow down the scientific progress of my project. | 348 | 1.32 | 1.5 | 0.82 | 0.02 |

*Note. N* = number of participants who answered the item; *M* = mean; *SD* = standard deviation.

### S7 Table: Univariate ANCOVAs and pairwise comparisons for the predictor Gender

| Summary ANCOVAs | | | | | |
|---|---|---|---|---|---|
| | Estimated marginal means | | | | |
| | Female | Male | Not Saying | *F*(2, 333) | $\eta^2$ |
| **Attitudes Scale** | -0.12 | -0.15 | -1.02 | 4.99** | .03 |
| **Subjective Norms Scale** | -1.04 | -1.03 | -1.36 | 0.39 | .00 |
| **Perceived Behavior Control – Resources Subscale** | -0.20 | -0.16 | -0.46 | 0.05 | .00 |
| **Perceived Behavior Control – Knowledge Subscale** | -0.35 | 0.10 | 0.89 | 16.35*** | .08 |
| **Intentions Scale** | 0.13 | -0.12 | -1.19 | 5.79** | .03 |
| **Motivations Scale** | -0.14 | -0.27 | -1.38 | 7.80*** | .04 |
| **Practical Obstacles Subscale** | 1.00 | 1.12 | 1.68 | 5.20** | .02 |
| **Competitive Obstacles Subscale** | 0.52 | 0.29 | 0.68 | 2.52 | .01 |

| Pairwise comparisons for significant ANCOVA effects | | | | | |
|---|---|---|---|---|---|
| | Estimated mean difference | *SE* | *df* | *t* | *p* |
| **Attitudes Scale** | | | | | |
| Female vs. Male | 0.03 | 0.13 | 356 | 0.24 | .969 |
| Female vs. Not saying | 0.89 | 0.33 | 356 | 2.72 | .019* |
| Male vs. Not saying | 0.86 | 0.32 | 356 | 2.68 | .021* |
| **Perceived Behavior Control – Knowledge Subscale** | | | | | |
| Female vs. Male | -0.45 | 0.13 | 337 | -3.38 | .002** |
| Female vs. Not saying | -1.24 | 0.35 | 337 | -3.50 | .002** |
| Male vs. Not saying | -0.79 | 0.35 | 337 | -2.25 | .064 |
| **Intentions Scale** | | | | | |
| Female vs. Male | 0.25 | 0.18 | 335 | 1.41 | .337 |
| Female vs. Not saying | 1.32 | 0.47 | 335 | 2.82 | .014* |
| Male vs. Not saying | 1.07 | 0.46 | 335 | 2.31 | .055 |
| **Motivations Scale** | | | | | |
| Female vs. Male | 0.12 | 0.14 | 334 | 0.85 | .672 |

| | | | | | |
|---|---|---|---|---|---|
| Female vs. Not saying | 1.23 | 0.38 | 334 | 3.29 | .003** |
| Male vs. Not saying | 1.11 | 0.37 | 334 | 3.01 | .008** |
| **Practical Obstacles Subscale** | | | | | |
| Female vs. Male | -0.12 | 0.12 | 334 | -1.03 | .561 |
| Female vs. Not saying | -0.68 | 0.29 | 334 | -2.34 | .052 |
| Male vs. Not saying | -0.56 | 0.29 | 334 | -1.97 | .122 |

*Note.* $F$ = $F$-statistic; $\eta^2$ = partial eta squared (effect size); *SE* = standard error; *df* = degrees of freedom; $t$ = $t$-statistic. Significant results are marked with $^{*}p < 0.05$, $^{**}p < 0.01$, $^{***}p < 0.001$. Tukey's correction was used for multiple comparisons.

### S8 Table: Univariate ANCOVAs and pairwise comparisons for the predictor Field of Research

**Summary ANCOVAs**

| | Estimated marginal means | | | | |
|---|---|---|---|---|---|
| | **Basic and experimental medical research** | **Basic biological research** | **General biology** | ***F*(2, 333)** | **$\eta^2$** |
| **Attitudes Scale** | -0.47 | -0.70 | -0.12 | 5.43** | .02 |
| **Subjective Norms Scale** | -1.16 | -1.13 | -1.14 | 0.13 | .00 |
| **Perceived behavioral Control – Resources Subscale** | -0.53 | -0.21 | -0.09 | 3.96* | .03 |
| **Perceived behavioral Control – Knowledge Subscale** | 0.13 | 0.23 | 0.28 | 0.43 | .01 |
| **Intentions Scale** | -0.46 | -0.69 | -0.02 | 2.60 | .02 |
| **Motivations Scale** | -0.61 | -1.03 | -0.15 | 7.02** | .04 |
| **Practical Obstacles Subscale** | 1.28 | 1.48 | 1.04 | 1.86 | .01 |
| **Competitive Obstacles Subscale** | 0.63 | 0.77 | 0.09 | 6.06* | .03 |

**Pairwise comparisons for significant ANCOVA effects**

| | **Estimated mean difference** | ***SE*** | ***df*** | ***t*** | ***p*** |
|---|---|---|---|---|---|
| **Attitudes Scale** | | | | | |
| Basic and experimental medical research vs. Basic biological research | 0.23 | 0.18 | 356 | 1.24 | .430 |
| Basic and experimental medical research vs. General biology | -0.35 | 0.19 | 356 | -1.91 | .136 |
| Basic biological research vs. General biology | -0.58 | 0.24 | 356 | -2.44 | .041* |
| **Perceived Behavioral Control – Resources Subscale** | | | | | |
| Basic and experimental medical research vs. Basic biological research | -0.32 | 0.17 | 337 | -1.87 | .149 |
| Basic and experimental medical research vs. General biology | -0.44 | 0.17 | 337 | -2.58 | .028* |
| Basic biological research vs. General biology | -0.12 | 0.22 | 337 | -0.54 | .850 |
| **Motivations Scale** | | | | | |
| Basic and experimental medical research vs. Basic biological research | 0.42 | 0.21 | 334 | 2.01 | .112 |
| Basic and experimental medical research vs. General biology | -0.47 | 0.21 | 334 | -2.22 | .070 |
| Basic biological research vs. General biology | -0.88 | 0.27 | 334 | -3.23 | .004** |

| | | | | | |
|---|---|---|---|---|---|
| **Competitive Obstacles Subscale** | | | | | |
| Basic and experimental medical research vs. Basic biological research | −0.14 | 0.17 | 334 | −0.79 | .707 |
| Basic and experimental medical research vs. General biology | 0.54 | 0.17 | 334 | 3.16 | .005** |
| Basic biological research vs. General biology | 0.68 | 0.22 | 334 | 3.04 | .007** |

*Note.* $F$ = $F$-statistic; $\eta^2$ = partial eta squared (effect size); *SE* = standard error; $df$ = degrees of freedom; $t$ = $t$-statistic.
Significant results are marked with *$p < 0.05$, **$p < 0.01$, ***$p < 0.001$. Tukey's correction was used for multiple comparisons.

**S9 Table: Univariate ANCOVAs and pairwise comparisons for the predictor Organization of Employment**

| **Summary ANCOVAs** | | | | | |
|---|---|---|---|---|---|
| | **Estimated marginal means** | | | | |
| | **Academic institution** | **Private industry** | **Other** | ***F*(2, 333)** | ***η²*** |
| **Attitudes Scale** | -0.64 | -0.30 | -0.36 | 2.40 | .01 |
| **Subjective Norms Scale** | -1.16 | -1.15 | -1.12 | 0.19 | .00 |
| **Perceived Behavior Control – Resources Subscale** | -0.26 | -0.28 | -0.28 | 0.04 | .00 |
| **Perceived Behavior Control – Knowledge Subscale** | 0.65 | -0.22 | 0.22 | 9.15*** | .05 |
| **Intentions Scale** | -0.49 | -0.31 | -0.38 | 0.45 | .00 |
| **Motivations Scale** | -0.68 | -0.69 | -0.41 | 0.48 | .00 |
| **Practical Obstacles Subscale** | 1.31 | 1.22 | 1.27 | 0.24 | .00 |
| **Competitive Obstacles Subscale** | 0.52 | 0.38 | 0.59 | 0.50 | .00 |
| **Pairwise comparisons for significant ANCOVA effects** | | | | | |
| | **Estimated mean difference** | ***SE*** | ***df*** | ***t*** | ***p*** |
| **Perceived Behavior Control – Knowledge Subscale** | | | | | |
| Academic institution vs. Private industry | 0.87 | 0.20 | 337 | 4.26 | <.001*** |
| Academic institution vs. Other | 0.43 | 0.24 | 337 | 1.78 | .177 |
| Private industry vs. Other | -0.43 | 0.30 | 337 | -1.44 | .321 |

*Note.* $\eta^2$ = partial eta squared (effect size); *SE* = standard error; *df* = degrees of freedom; *t* = *t*-statistic.
Significant results are marked with *$p < 0.05$, **$p < 0.01$, ***$p < 0.001$. Tukey's correction was used for multiple comparisons.

### S10 Table 1: Barriers

| Open-ended items |
|---|
| *All participants* |
| ***What do you perceive as drawbacks of preregistration?*** |
| Number of responses = 142 |
| ***What do you think would be the long-term negative consequences of mandatory preregistration?*** |
| Number of responses = 140 |
| *Only if participants never preregistered* |
| ***What are the reasons for not preregistering your studies?*** |
| Number of responses = 313 |
| *Only if participants preregistered at least one study* |
| ***I am now less motivated to preregister than I was before because…*** |
| Number of responses = 5 |

### S10 Table 2: Facilitators

| Open-ended items |
|---|
| *All participants* |
| ***What do you perceive as benefits of preregistration?*** |
| Number of responses = 140 |
| ***What do you think would be the long-term benefits of mandatory preregistration?*** |
| Number of responses = 140 |
| *Only if participants preregistered at least one study* |
| ***I am now more motivated to preregister than I was before because…*** |
| Number of responses = 6 |

### S10 Table 3: Suggestions for improvement

| Open-ended items | |
|---|---|
| | *All participants* |
| ***We are interested in how we can improve various aspects of preregistration (e.g. templates, repositories, reviewing process, integrations in published articles, education etc.). Do you have any suggestions? What do you think should be improved about preregistration?*** | |
| Number of responses = 129 | |
| ***What would make you (and perhaps other researchers) preregister more often?*** | |
| Number of responses = 137 | |
| ***Do you have any suggestions as to how to lower your (or other researchers') negative perceptions of preregistration?*** | |
| Number of responses = 131 | |
| **Close-ended items** | |
| ***The application form for animal experiments (Animex-ch) and the preregistration template should be the same.*** | |
| Disagree | 24.9% (85) |
| Neutral | 28.7% (98) |
| Agree | 46.5% (159) |
| *N* | 342 (76 missing) |

| | |
|---|---|
| ***The application form for animal experiments (Animex-ch) and the preregistration template should be separate from each other.*** | |
| Disagree | 39.8% (137) |
| Neutral | 31.1% (107) |
| Agree | 29.1% (100) |
| *N* | 344 (74 missing) |
| ***Certain sections of the application form for animal experiments (Animex-ch) should be linked and uploaded to the preregistration template.*** | |
| Disagree | 21.0% (72) |
| Neutral | 28.6% (98) |
| Agree | 50.4% (173) |
| *N* | 343 (75 missing) |

*Note. N* = total sample size
